# Neuronal origins of biases in economic choices under sequential offers

**DOI:** 10.1101/2021.11.07.467645

**Authors:** Weikang Shi, Sébastien Ballesta, Camillo Padoa-Schioppa

## Abstract

Economic choices are characterized by a variety of biases. Understanding their origins is a long-term goal for neuroeconomics, but progress on this front has been limited. Here we examined choice biases observed when two goods are offered sequentially. In the experiments, rhesus monkeys chose between different juices offered simultaneously or in sequence. Choices under sequential offers were less accurate (higher variability). They were also biased in favor of the second offer (order bias) and in favor of the preferred juice (preference bias). Analysis of neuronal activity recorded in orbitofrontal cortex revealed that these phenomena emerged at different computational stages. The lower choice accuracy reflected weaker offer value signals (valuation stage), the order bias emerged during value comparison (decision stage), and the preference bias emerged late in the trial (post-comparison). Our approach, leveraging recent notions on the neural mechanisms of economic decisions, may shed light on other aspects of choice behavior.

## Introduction

Some of the most mysterious aspects of economic behavior are choice biases documented in behavioral economics (Camerer et al., 2003; Kahneman and Tversky, 2000; Lichtenstein and Slovic, 2006). Standard economic theory fails to account for these effects, and shedding light on their origins is a long-term goal for neuroeconomics (Camerer et al., 2005; Glimcher and Rustichini, 2004). Progress on this front has been relatively modest, largely because the neural underpinnings of (even simple) choices were poorly understood until recently. However, the last 15 years have witnessed substantial advances. An important turning point was the development of experimental protocols in which subjects choose between different goods and relative subjective values are inferred from choices. Decision variables defined from these values are used to interpret neural activity (Kable and Glimcher, 2007; Padoa-Schioppa and Assad, 2006; Plassmann et al., 2007). Studies that adopted this paradigm showed that neurons in numerous brain regions represent the values of offered and chosen goods (Amemori and Graybiel, 2012; Cai et al., 2011; Cai and Padoa-Schioppa, 2012; Hosokawa et al., 2013; Jezzini and Padoa-Schioppa, 2020; Kim et al., 2008; Lak et al., 2014; Levy et al., 2010; Louie and Glimcher, 2010; Padoa-Schioppa and Assad, 2006; Pastor-Bernier et al., 2019; Shenhav and Greene, 2010). Furthermore, recent experiments using electrical stimulation showed that offer values encoded in the orbitofrontal cortex (OFC) are causally linked to choices (Ballesta et al., 2020). These results are of high significance for three reasons.

First, the identification in OFC and other brain regions of distinct groups of neurons encoding different decision variables is essential to ultimately understand the neural circuit and the mechanisms through which economic decisions are formed.

Second, in a more conceptual sense, the results summarized above provide a long-sought validation for the construct of value. The proposal that choices entail computing and comparing subjective values was put forth by early economists such as Bernoulli and Bentham (Niehans, 1990). Although this idea has remained influential, values defined at the behavioral level suffer from a fundamental problem of circularity. On the one hand, choices supposedly maximize values; on the other hand, values cannot be measured behaviorally independent of choices (Samuelson, 1938). Because of this problem, the construct of value gradually lost centrality in economic theory. Thus in the standard neoclassic formulation choices are “as if” driven by values, but there is no commitment to the idea that agents actually compute values (Samuelson, 1947). In this perspective, the fact that neuronal firing rates in any brain region are linearly related to values defined at the behavioral level constitutes powerful evidence that choices indeed entail the computation of values (Camerer, 2008).

Third and less frequently discussed, the identification of neurons encoding offer values and other decision variables, together with some rudimentary understanding of the decision circuit, provides the opportunity to break the circularity problem described above. To appreciate this point, consider the fact that economic choices are often affected by seemingly idiosyncratic biases. For example, while choosing between two options offered sequentially, people and monkeys typically show a bias favoring the second option (Ballesta and Padoa-Schioppa, 2019; Krajbich et al., 2010; Rustichini et al., 2021). This order bias might occur for at least two reasons. (1) Subjects might assign a higher value to any given good if that good is offered second. (2) Alternatively, subjects might assign identical values independent of the presentation order, and the bias might emerge downstream of valuation, for example during value comparison. In the latter scenario, by introducing the order bias, the decision process would actually fail to maximize the value obtained by the agent. Due to the circularity problem described above, these two hypotheses are ultimately not distinguishable based on behavior alone. However, access to a credible neural measure for the offer values makes it possible, at least in principle, to disambiguate between them. The results presented in this study build on this fundamental idea.

We focused on choice biases measured when two goods are offered sequentially. In the experiments, monkeys chose between two juices offered in variable amounts. In each session, we randomly interleaved two types of trials referred as two tasks. In Task 1, offers were presented simultaneously; in Task 2, offers were presented in sequence. Comparing choices across tasks revealed three phenomena. (1) Monkeys were substantially less accurate (higher choice variability) in Task 2 (sequential offers) compared to Task 1 (simultaneous offers). (2) Choices in Task 2 were biased in favor of the second offer (order bias). (3) Choices in Task 2 were biased in favor of the preferred juice (preference bias) (Shi et al., 2021). These effects are especially interesting because in most daily situations offers available for choice appear or are examined sequentially. In the present study, we investigated the neuronal origins of these phenomena, collectively referred to as choice biases.

Neuronal recordings focused on the OFC. Earlier work on choices under simultaneous offers identified in this area different groups of cells encoding individual offer values, the binary choice outcome (chosen juice), and the chosen value (Padoa-Schioppa, 2013; Padoa-Schioppa and Assad, 2006). Furthermore, previous analyses indicated that choices under sequential offers engage the same neuronal populations (Ballesta and Padoa-Schioppa, 2019; Shi et al., 2021). In other words, the cell groups labeled *offer value*, *chosen juice* and *chosen value* can be identified in either choice task and appear to preserve their functional role. In first approximation, the variables encoded in OFC capture both the input (offer values) and the output (chosen juice, chosen value) of the choice process, suggesting that the cell groups identified in this area constitute the building blocks of a decision circuit (Padoa-Schioppa and Conen, 2017). A series of experimental (Ballesta et al., 2020; Camille et al., 2011; Rich and Wallis, 2016) and theoretical(Friedrich and Lengyel, 2016; Rustichini and Padoa-Schioppa, 2015; Solway and Botvinick, 2012; Song et al., 2017; Zhang et al., 2018) results support this view. Here we put forth a more articulated computational framework. In our account, different groups of OFC neurons participate in value computation and value comparison, and these processes are embedded in an ensemble of mental operations taking place before, during and after the decision itself. In this view, sensory information, memory traces and internal states are processed upstream of OFC and integrated in the activity of *offer value* cells. These neurons provide the primary input to a circuit formed by *chosen juice* cells and *chosen value* cells, where values are compared. The output of this circuit feeds brain regions involved in working memory and the construction of action plans (**Fig.1**).

**Figure 1.**
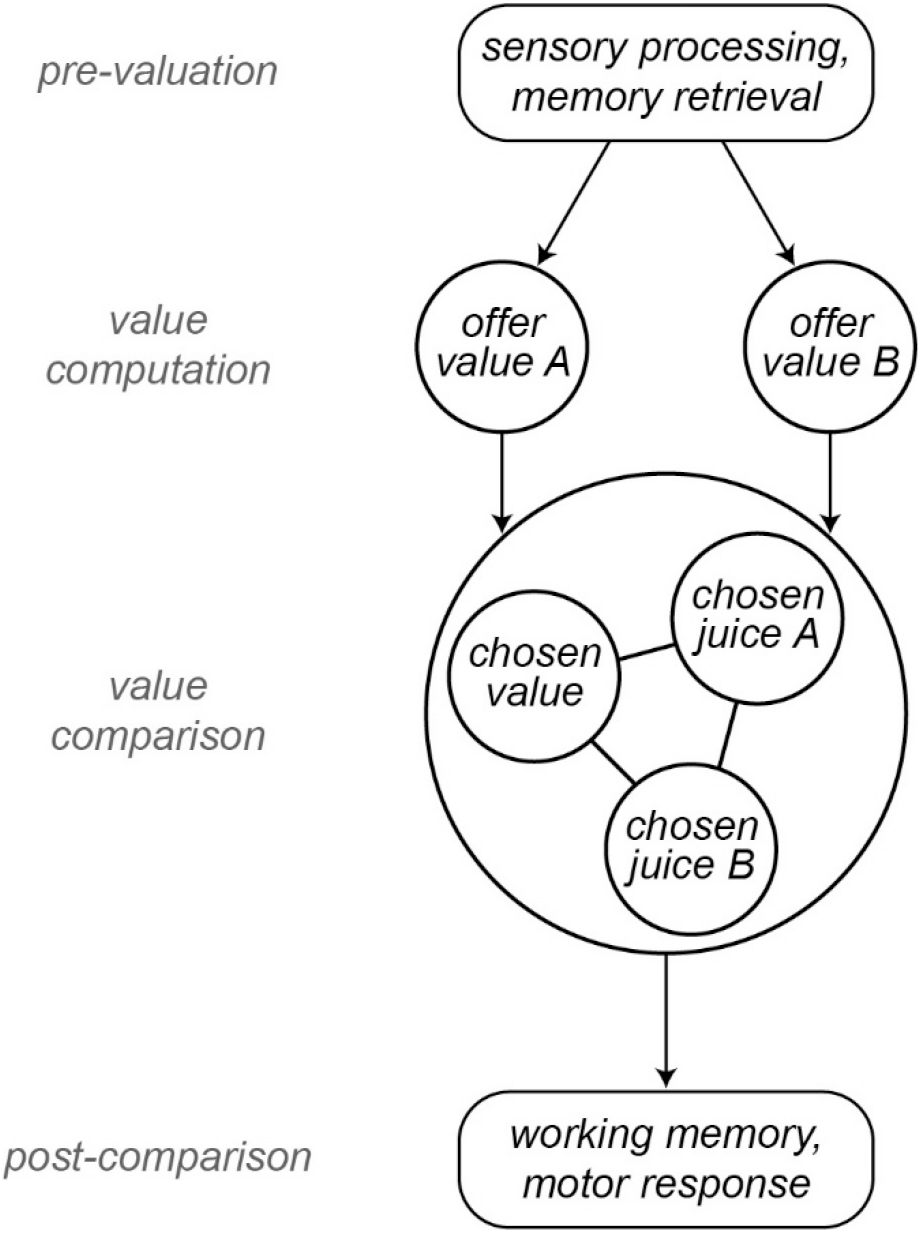
Computational framework. Information about sensory input, stored memory and the motivational state is integrated during the computation of offer values. In OFC, *offer value* cells provide the primary input to a decision circuit composed of *chosen juice* cells and *chosen value* cells. The detailed structure of the decision circuit is not well understood, but previous work indicates that decisions under sequential offers rely on circuit inhibition. In essence, neurons encoding the value of the first offer (offer1) indirectly impose a negative offset on the activity of *chosen juice* cells associated with the second offer (offer2). Notably, this circuit might also subserve working memory. The decision output, captured by the activity of *chosen juice* cells, informs other brain regions that maintain it in working memory and transform it into a suitable action plan. Choice measured behaviorally is ultimately defined by the motor response. This framework highlights the fact that choice biases and/or noise might emerge at multiple computational stages. The arrows indicated here capture only the primary connections.

This framework guided a series of analyses relating the activity of each cell group to the choice biases described above. Our results revealed that different phenomena emerged at different computational stages. The lower choice accuracy observed under sequential offers reflected weaker offer value signals (valuation stage). Conversely, the order bias did not have neural correlates at the valuation stage, but rather emerged during value comparison (decision stage). Finally, the preference bias did not have neural correlates at the valuation stage or during value comparison; it emerged late in the trial, shortly before the motor response.

## Results

### Choice biases under sequential offers

Two monkeys participated in the experiments. In each session, they chose between two juices labeled A and B, with A preferred. Offers were represented by sets of colored squares on a monitor, and animals indicated their choice with a saccade. In each session, two choice tasks were randomly interleaved. In Task 1, offers were presented simultaneously (**Fig.2A**); in Task 2, offers were presented in sequence (**Fig.2B**). A cue displayed at the beginning of the trial revealed to the animal the task for that trial. Offers varied from trial to trial, and we indicate the quantities offered in any given trial with *q_A_* and *q_B_*. An “offer type” was defined by two quantities [*q_A_*, *q_B_*], and the same offer types were used for the two tasks in each session. For Task 2, trials in which juice A was offered first and trials in which juice B was offered first are referred to as “AB trials” and “BA trials”, respectively. The first and second offers are referred to as “offer1” and “offer2”, respectively.

**Figure 2.**
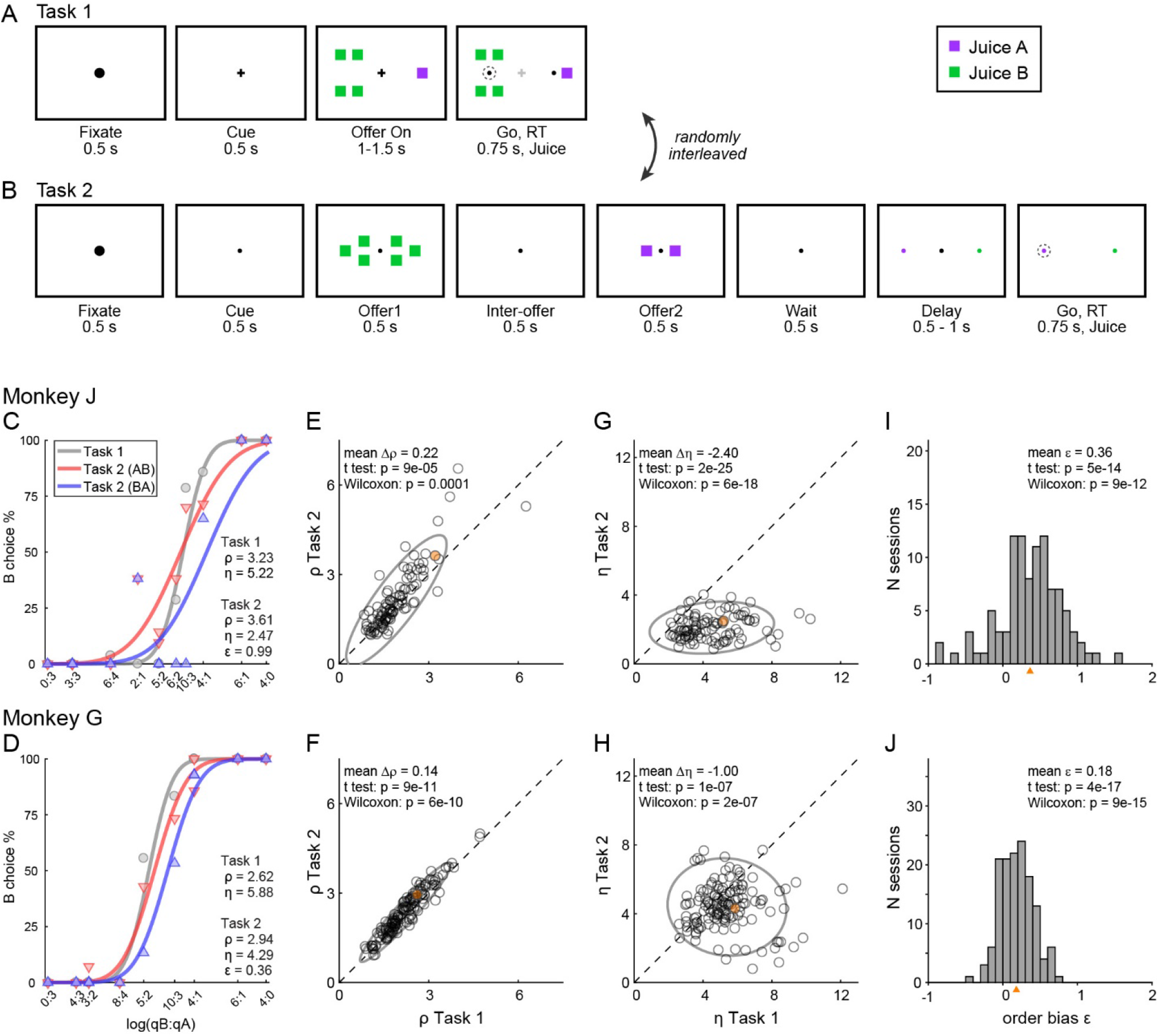
Experimental design and choice biases. **(AB)** Experimental design. Animals chose between two juices offered in variable amounts. Offers were represented by sets of color squares. For each offer, the color indicated the juice type and the number of squares indicated the juice amount. In each session, trials with Task 1 and Task 2 were randomly interleaved. In Task 1, two offers appeared simultaneously on the left and right sides of the fixation point. In Task 2, offers were presented sequentially, spaced by an inter-offer delay. After a wait period, two saccade targets matching the colors of the offers appeared on the two sides of the fixation point. The left/right configuration in Task 1, the presentation order in Task 2 and the left/right position of the saccade targets in Task 2 varied randomly from trial to trial. In any session, the same set of offer types was used for both tasks. **(C)** Example session 1. The percent of B choices (y-axis) is plotted against the log quantity ratio (x-axis). Each data point indicates one offer type in Task 1 (gray circles) or Task 2 (red and blue triangles for AB trials and BA trials, respectively). Sigmoids were obtained from probit regressions. The relative value (ρ) and sigmoid steepness (*η*) measured in each task and the order bias (*ε*) measured in Task 2 are indicated. In this session, the animal presented all three biases. Compared to Task 1, choices in Task 2 were less accurate (*η_Task2_* < *η_Task1_*) and biased in favor of juice A (*ρ_Task2_* > *ρ_Task1_*; preference bias). Furthermore, choices in Task 2 were biased in favor of offer2 (*ε* > 0; order bias). **(D)** Example session 2. Same format as panel C. **(EF)** Comparing relative value across choice tasks. Each data point represents one session and gray ellipses indicate 90% confidence intervals. For both monkeys, relative values in Task 2 (y-axis) were significantly higher than in Task 1 (x-axis). Furthermore, the main axis of each ellipse was rotated counterclockwise compared to the identity line. **(GH)** Comparing the sigmoid steepness across choice tasks. For both monkeys, sigmoids were consistently shallower (smaller *η*) in Task 2 compared to Task 1. **(IJ)** Order bias, distribution across sessions. Both monkeys presented a consistent bias favoring offer2 (mean(*ε*)>0). Panels CEGI are from monkey J (N = 101 sessions); panels DFHJ are from monkey G (N = 140 sessions). Sessions shown in panels CD are highlighted in yellow in panels EFGH. Triangles in panels IJ indicate the mean. Statistical tests and exact p values are indicated in each panel.

The data set included 241 sessions (101 from monkey J, 140 from monkey G; see **Methods**). Sessions lasted for 217-880 trials (mean ± std = 589 ± 160). For each session, we analyzed choices in the two tasks separately using probit regressions. For Task 1 (simultaneous offers), we used the following model:

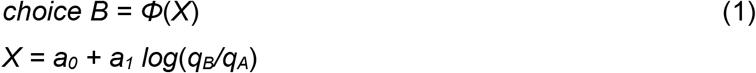

where *choice B* = 1 if the animal chose juice B and 0 otherwise, *Φ* was the cumulative function of the standard normal distribution, and *q_A_* and *q_B_* were the quantities of juices offered on any given trial. From the fitted parameters *a_0_* and *a_1_*, we derived measures for the relative value of the two juices *ρ_Task1_* = *exp*(−*a_0_/a_1_*) and for the sigmoid steepness *η_Task1_* = *a_1_*. Intuitively, the relative value was the quantity ratio *q_B_*/*q_A_* that made the animal indifferent between the two juices, and the sigmoid steepness was inversely related to choice variability.

For Task 2 (sequential offers), we used the following model:

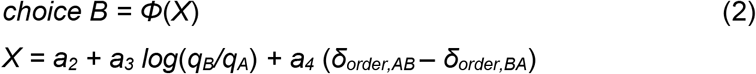

where *δ_order,AB_* = 1 for AB trials and 0 otherwise, and *δ_order,BA_* = 1 – *δ_order,AB_*. In essence, AB trials and BA trials were analyzed separately but assuming that the two sigmoids were parallel. From the fitted parameters *a_2_*, *a_3_* and *a_4_*, we derived measures for the relative value of the two juices *ρ_Task2_* = *exp*(−*a_2_/a_3_*), for the sigmoid steepness *η_Task2_* = *a_3_*, and for the order bias *ε* = *2 ρ_Task2_ a_4_/a_3_*. Intuitively, the order bias was a bias favoring the first or the second offer. Specifically, *ε*<0 indicated a bias favoring offer1; *ε*>0 indicated a bias favoring offer2. We also defined relative values specific to AB trials and BA trials as *ρ_AB_* = *exp*(−(*a_2_*+*a_4_*)/*a_3_*) and *ρ_BA_* = *exp*(−(*a_2_*-*a_4_*)/*a_3_*). Of note, the order bias was defined such that

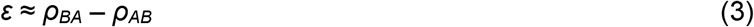

The experimental design gave us the opportunity to compare choices across tasks independently of factors such as selective satiation or changes in the internal state. The relative values measured in the two tasks were highly correlated (**Fig.2EF**). At the same time, our analyses revealed three interesting phenomena. First, for both animals, sigmoids measured in Task 2 were significantly shallower compared to Task 1 (**Fig.2GH**). In other words, presenting offers in sequence reduced choice accuracy. Second, in Task 2, both animals showed a consistent order bias favoring offer2 (**Fig.2IJ**). Third, in both animals, relative values in Task 2 were significantly higher than in Task 1 (*ρ_Task2_*>*ρ_Task1_*), and this effect increased with the relative value (**Fig.2EF**). In other words, the ellipse marking the 90% confidence interval for the joint distribution of relative values laid above the identity line and was rotated counterclockwise compared to the identity line.

To further investigate the differences in relative values measured across tasks, we quantified them separately in AB trials and BA trials in each monkey. We thus examined the relation between *ρ_Task1_* and *ρ_Task2,AB_* and, separately, that between *ρ_Task1_* and *ρ_Task2,BA_* (**Fig.3**). In both animals and in both sets of trials, the ellipse marking the 90% confidence interval was rotated counterclockwise compared to the identity line. Furthermore, the ellipse measured for BA trials was higher than that for AB trials. We quantified these observations with an analysis of covariance (ANCOVA) using the presentation order (AB, BA) as a covariate and imposing parallel lines (**Fig.3C,F**). In both animals, the two regression lines were significantly distinct (difference in intercept >0, p ≤ 0.002 in each animal). This result confirmed the presence of an order bias favoring offer2 in Task 2. Concurrently, in both animals the regression slope was significantly >1 (p ≤ 0.04 in each animal; ellipse rotation). This result indicated that the animals had an additional bias favoring juice A in Task 2, and that this bias increased as a function of the relative value *ρ*. We refer to this phenomenon as the preference bias.

**Figure 3.**
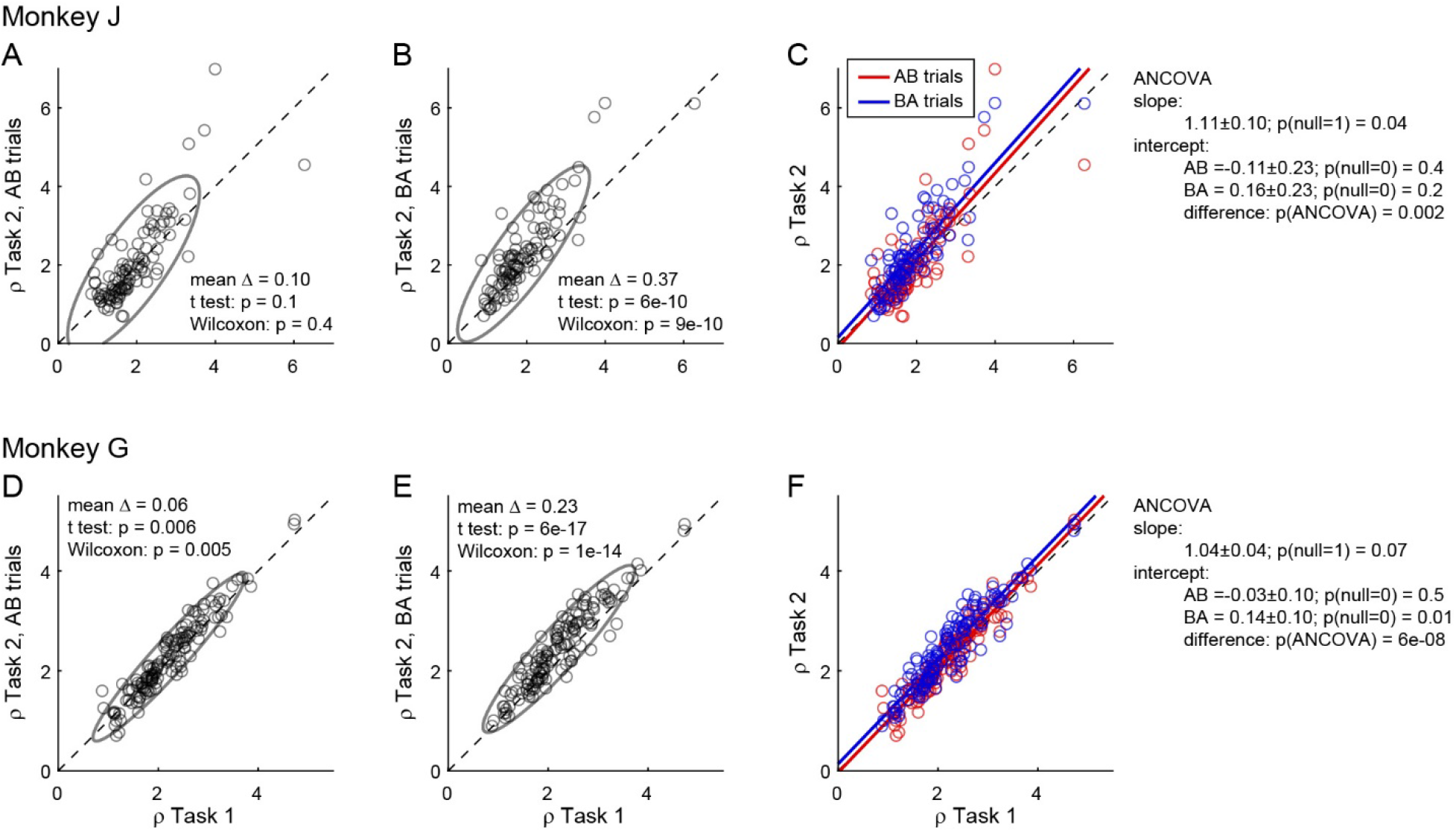
Order bias and preference bias. **(ABC)** Monkey J (N = 101 sessions). In panels A and B, *ρ_Task2,AB_* and *ρ_Task2,BA_* (y-axis) are plotted against *ρ_Task1_* (x-axis). Each data point represents one session and gray ellipses indicate 90% confidence intervals. The main axis of both ellipses is rotated counterclockwise compared to the identity line (preference bias). In addition, the ellipse in panel B is displaced upwards compared to that in panel A (order bias). In panel C, the same data are pooled and color coded. The two lines are from an ANCOVA (covariate: order; parallel lines). The regression slope is significantly >1 (preference bias) and the two intercepts differ significantly from each other (order bias). **(DEF)** Monkey G (N = 140 sessions). Same format. The results closely resemble those for monkey J but the preference bias is weaker.

### Origins of choice biases: Computational framework

The following sections present a series of results on the neuronal origins of these biases. We begin by discussing the computational framework for these analyses.

Economic choice is thought to entail two stages: values are assigned to the available offers and a decision is made by comparing values. Importantly, in our tasks and in most circumstances, choices elicit an ensemble of mental operations taking place before, during and after the computation and comparison of offer values. Upstream of valuation, choices examined here entail the sensory processing of visual stimuli and the retrieval from memory of relevant information (e.g., the association between color and juice type). Downstream of value comparison, the decision outcome must guide a suitable motor response. In addition, performance in Task 2 requires holding in working memory the value of offer1 until offer2, remembering the decision outcome for an additional delay, and mapping that outcome onto the appropriate saccade target (**Fig.2B**). In principle, choice biases could emerge at any of these computational stages. Likewise, each of these mental operations could be noisy and thus contribute to choice variability.

Neuronal activity in OFC does not capture all of these processes. However, previous work indicates that neurons in this area participate both in value computation and value comparison. In the framework proposed here (**Fig.1**), sensory and limbic areas feed *offer value* cells, where values are integrated. In turn, *offer value* cells provide the primary input to a neural circuit constituted by *chosen juice* cells and *chosen value* cells, where decisions are formed. Finally, the decision circuit is connected with downstream areas, such as lateral prefrontal cortex, engaged in working memory and in transforming choice outcomes into suitable action plans. This scheme reflects the anatomical connectivity of OFC and other prefrontal regions (Carmichael and Price, 1995a, b; Petrides and Pandya, 2006; Saleem et al., 2013; Takahara et al., 2012); it is motivated by neurophysiology results from OFC (Ballesta et al., 2020; Rich and Wallis, 2016) and connected areas (Cai and Padoa-Schioppa, 2014; Sasikumar et al., 2018); and it is consistent with computational models of economic decisions (Friedrich and Lengyel, 2016; Rustichini and Padoa-Schioppa, 2015; Solway and Botvinick, 2012; Song et al., 2017; Yim et al., 2019; Zhang et al., 2018).

Of note, both *offer value* and *chosen value* cells encode subjective values. However, in the framework of **Fig.1**, *offer value* cells express a pre-decision value, while *chosen value* cells express a value emerging during the decision process. Conversely, the activity of *chosen juice* cells captures the evolving commitment to a particular choice outcome. In this framework, suitable analyses of neuronal activity may reveal whether particular choice biases emerge during valuation, during value comparison, or in subsequent computational stages.

### Reduced accuracy under sequential offers emerged at the valuation stage

Other things equal, choices under sequential offers (Task 2) were significantly less accurate than choices under simultaneous offers (Task 1; **Fig.2**). We first investigated the neural origins of this phenomenon.

The primary data set examined in this study included 183 *offer value* cells, 160 *chosen juice* cells and 174 *chosen value* cells (see **Methods**). Comparing neuronal responses across tasks, we noted that offer value signals in Task 2 were significantly weaker than in Task 1. **Fig.4AC** illustrates one example cell. In both tasks, this neuron encoded the *offer value B*. However, the activity range (see **Methods**) measured in Task 2 was smaller than that measured in Task 1. This effect was also observed at the population level. For this analysis, we pooled *offer value* cells associated with juices A and B, and with positive or negative encoding (see **Methods**). For Task 1, we focused on the post-offer time window; for Task 2, we focused on post-offer1 and post-offer2 time windows, pooling trial types from both windows. For each cell, we imposed that the response be significantly tuned in these time windows in each task, and we quantified the mean activity and the activity range (*Δr*, see **Methods**). At the population level, the mean activity did not differ significantly across tasks (p = 0.6, t test; p = 0.4, Wilcoxon test **Fig.4D**). In contrast, the activity range was significantly lower in Task 2 compared to Task 1 (*Δr_Task2_* < *Δr_Task1_*; p = 0.06, t test; p = 0.02, Wilcoxon test **Fig.4E**). In other words, offer value signals were weaker in Task 2 compared to Task 1.

**Figure 4.**
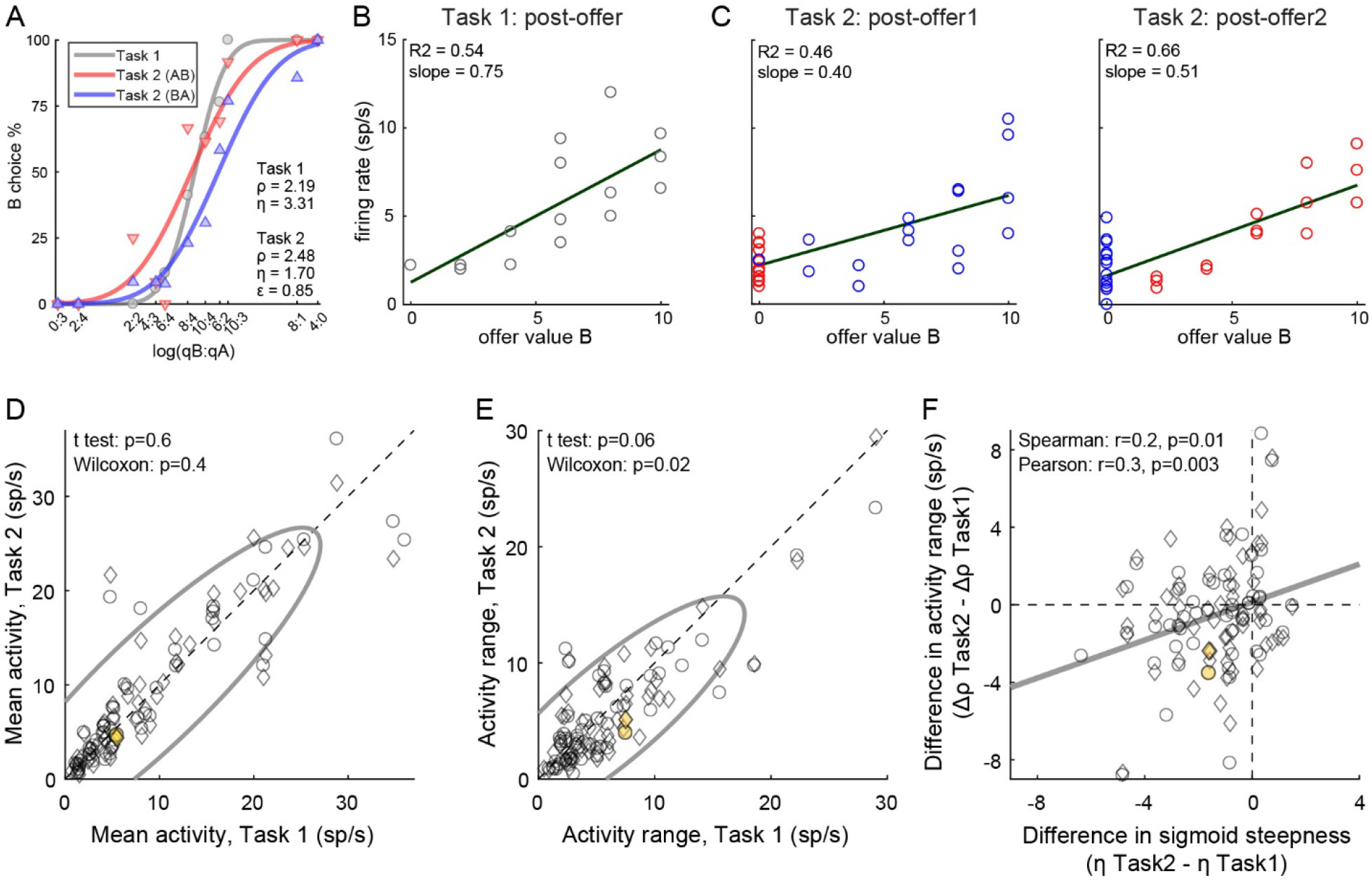
Lower choice accuracy in Task 2 reflects weaker offer value signals. **(A-C)** Weaker offer value signals in Task 2, example cell. Panel A illustrates the choice pattern. Panel B illustrates the neuronal response measured in Task 1 (post-offer time window). Each data point represents one trial type. In C, two panels illustrate the neuronal responses measured in Task 2 (post-offer1 and post-offer2 time windows). Each data point represents one trial type; red and blue colors are for AB and BA trials, respectively. In panels B and C, firing rates (y-axis) are plotted against variable *offer value B* and gray lines are from linear regressions. Notably, the cell has lower activity range in Task 2 than in Task 1. **(DE)** Weaker offer value signals in Task 2, population analysis (N = 109 *offer value* cells). The two panels illustrate the results for the mean activity and the activity range, respectively. In each panel, x-axis and y-axis represent measures obtained in Task 1 and Task 2, respectively. Each data point represents one cell. For each cell, we examined one time window (post-offer) in Task 1 and two time windows (post-offer1 and post-offer2) in Task 2. Circles and diamonds refer to post-offer1 and post-offer2 time windows, respectively. Measures of mean activity measured in the two tasks (panel D) were statistically indistinguishable. In contrast, activity ranges (panel E) were significantly reduced in Task 2 compared to Task 1. Statistical tests and exact p values are indicated in each panel. The example cell shown in panels A-C is highlighted in orange in panels DE. **(F)** Offer value signals and choice accuracy (N = 109 cells). For each *offer value* cell, we computed the activity range *Δr* in each task (see **Methods**). Here the difference in activity range *ΔΔr* = *Δr_Task2_* – *Δr_Task1_* (y-axis) is plotted against the difference in sigmoid steepness *Δη* = *η_Task2_* – *η_Task1_* measured in the same session (x-axis). The two measures were significantly correlated across the population. The gray line in panel F is from a linear regression. This analysis was restricted to 53 cells significantly tuned in the post offer time window (Task 1) and post offer1 time window (Task 2), and 56 cells significantly tuned in the post offer time window (Task 1) and post offer2 time window (Task 2).

The activity of *offer value* cells is causally related to choices (Ballesta et al., 2020). Furthermore, for given value range and mean activity, the activity range determines the neuronal signal-to-noise ratio. Indeed, we previously found that decreases in the encoding slope of *offer value* cells due to range adaptation reduce choice accuracy (Conen and Padoa-Schioppa, 2019; Rustichini et al., 2017). Along similar lines, we inquired whether the difference in choice accuracy measured across tasks (**Fig.2GH**) might be explained, at last partly, by differences in neuronal activity range (**Fig.4E**). We thus examined the relation between the difference in sigmoid steepness (*Δη* = *η_Task2_* – *η_Task1_*) and the difference in activity range (*ΔΔr* = *Δr_Task2_* – *Δr_Task1_*). The two measures were positively correlated (Spearman r = 0.2, p = 0.01; Pearson r = 0.3, p = 0.003; **Fig.4F**). In other words, the drop in choice accuracy observed in Task 2 compared to Task 1 correlated with weaker offer value signals. Of note, similar analyses on *chosen value* cells and *chosen juice* cells yielded negative results (**Fig.S1**).

In conclusion, the lower choice accuracy measured in Task 2 compared to Task 1 correlated with weaker offer value signals in OFC. Thus this behavioral phenomenon emerged, at least partly, during valuation.

### The order bias emerged during value comparison

The next series of analyses focused on the neural origins of the order bias. Since this phenomenon pertains only to choices under sequential offers, we included in the analyses an additional data set recorded in the same animals performing only Task 2 (see **Methods**).

In the framework of **Fig.1**, we first inquired whether the order bias emerged during valuation. If this was the case, for any given good, *offer value* cells should encode a higher value when the good is presented as offer2. To test this hypothesis, we pooled *offer value* cells associated with the two juices. For each neuron, ‘E’ indicated the juice encoded by the cell and ‘O’ indicated the other juice. We thus refer to EO trials and OE trials. For any given cell, we compared the response recorded in EO trials (post-offer1 time window) with the response recorded in OE trials (post-offer2 time window). If the order bias emerged during valuation, the tuning intercept and/or the tuning slope should be higher for the latter (**Fig.S2A**). Contrary to this prediction, across a population of 128 cells, we did not find any systematic difference in intercept or slope (**Fig.S2BC**). Furthermore, the difference between the intercepts and slopes measured in OE and EO trials did not correlate with the order bias (**Fig.S2D**). In conclusion, assigned values did not depend on the presentation order.

We next examined whether the order bias emerged during value comparison. If so, the bias should be reflected in the activity of both *chosen juice* and *chosen value* cells (**Fig.1**). For *chosen value* cells, the hypothesis might be tested noting that in post-offer1 and post-offer2 time windows these neurons encoded the value currently offered independently of the juice type (**Table S1**). Thus the activity measured in these time windows in AB and BA trials provided neuronal measures for the relative values of the two juices. More specifically, for each *chosen value* cell, we derived the two measures *ρ^neuronal^_AB_* and *ρ^neuronal^_BA_* for AB trials and BA trials, respectively (**Fig.5A**; see **Methods**). We also defined the difference *Δρ^neuronal^* = *ρ^neuronal^_BA_* – *ρ^neuronal^_AB_*. We recall that the order bias was essentially equal to the difference between the relative values measured behaviorally in BA and AB trials (**Eq.3**). Thus, assessing whether the activity of *chosen value* cells reflected the order bias amounts to testing the relation between *Δρ^neuronal^* and *ε*.

**Figure 5.**
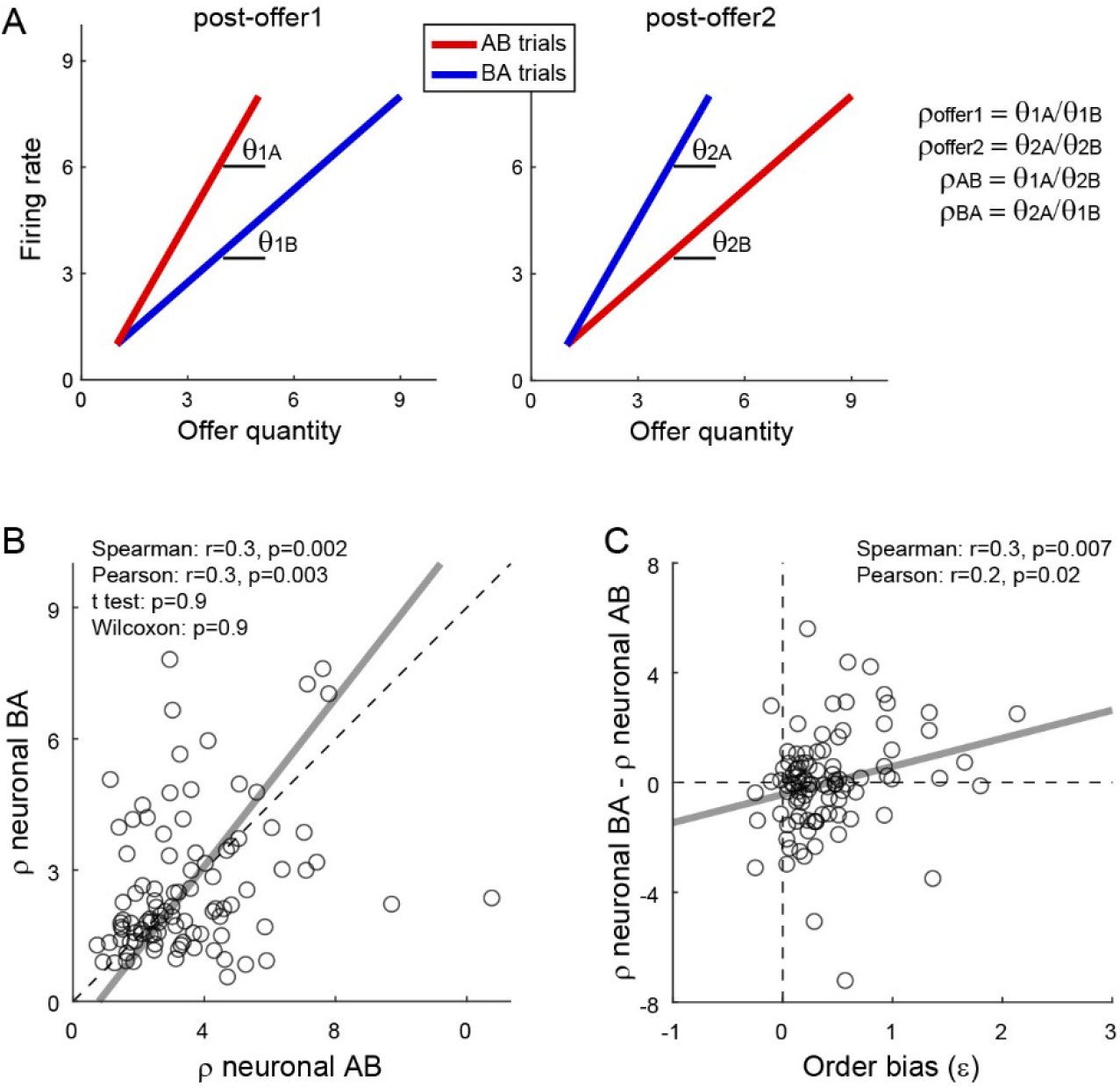
Fluctuations in order bias and fluctuations in the activity of *chosen value* cells. **(A)** Neuronal measures of relative value. The two panels represent in cartoon format the response of a *chosen value* cell in the post-offer1 and post-offer2 time window (Task 2). In each of these time windows, *chosen value* cells encode the value of the offer on display. Here the two axes correspond to the firing rate (y-axis) and to the offered juice quantity (x-axis). The two colors correspond to the two orders (AB, BA). In each time window, two linear regressions provide two slopes, proportional to the value of the two juices. From the four measures *θ_1A_* (left panel, red), *θ_1B_* (left panel, blue), *θ_2A_* (right panel, blue) and *θ_2B_* (right panel, red), we derive four neuronal measures of relative value (**Methods, Eqs.9-12**). **(B)** Neuronal measures of relative value in AB trials and BA trials (N = 96 cells). The x- and y-axis correspond to *ρ^neuronal^_AB_* and *ρ^neuronal^_BA_*, respectively. Each data point represents one cell. The two measures are strongly correlated. The gray line is from a Deming regression. **(C)** Fluctuations of relative value and fluctuations in order bias (N = 96 cells). For each *chosen value* cell, we quantified the difference in the neuronal measure of relative value *Δρ^neuronal^* = *ρ^neuronal^_AB_* – *ρ^neuronal^_BA_*. Here, the x-axis is the order bias (*ε*), the y-axis is *Δρ^neuronal^*, and each data point corresponds to one cell. Although *Δρ^neuronal^* was on average close to 0 (panel B), fluctuations of *Δρ^neuronal^* correlated with fluctuations of *ε* across the population. The gray line is from a linear regression. Statistical tests and exact p values are indicated in each panel. This analysis was restricted to 96 cells that had significant *θ_1A_*, *θ_1b_*, *θ_2A_* and *θ_2B_*.

We conducted a population analysis of 96 *chosen value* cells. Confirming previous results (Padoa-Schioppa and Assad, 2006), neuronal and behavioral measures of relative value were highly correlated. The two neuronal measures or relative value, *ρ^neuronal^_AB_* and *ρ^neuronal^_BA_*, did not differ significantly from each other (**Fig.5B**). However, and most importantly, the difference *Δρ^neuronal^* and the order bias *ε* were significantly correlated across the population (Spearman r = 0.3, p = 0.007; Pearson r = 0.2, p = 0.02; **Fig.5C**). Hence, session-to-session fluctuations in the activity of *chosen value* cells correlated with fluctuations in the order bias.

Further insights on the order bias came from the analysis of *chosen juice* cells. Again, for each neuron, E and O indicated the juice encoded by the cell and the other juice, respectively. A previous study found that the baseline activity of *chosen juice* cells recorded in OE trials immediately before offer2 was negatively correlated with the value of offer1 (i.e., the value of the other juice) – a phenomenon termed circuit inhibition (Ballesta and Padoa-Schioppa, 2019). If the decision is conceptualized as the evolution of a dynamic system (Rustichini and Padoa-Schioppa, 2015; Wang, 2002), circuit inhibition sets the system’s initial conditions and is thus integral to value comparison. In this account, the evolving decision is essentially captured by the activity of *chosen juice* cells in OE trials, which reflects a competition between the negative offset set by the value of offer1 (initial condition) and the incoming signal encoding the value of offer2. If so, the intensity of circuit inhibition should be negatively correlated with the order bias.

We tested this prediction as follows. First, we replicated previous findings and confirmed the presence of circuit inhibition in our primary data set (**Fig.6A**). Second, we focused on a 300 ms time window starting 250 ms before offer2 onset. For each *chosen juice* cell, we regressed the firing rate against the normalized offer1 value (see **Methods**). Thus the regression slope *c_1_* quantified circuit inhibition for individual cells. Across a population of 295 *chosen juice* cells, mean(*c_1_*) was significantly <0 (p = 5 10^-6^, t test; p = 9 10^-8^, Wilcoxon test; **Fig.6B**). Third, we examined the relation between circuit inhibition (*c_1_*) and the order bias (*ε*). Confirming the prediction, the two measures were significantly correlated across the population (Spearman r = 0.1, p = 0.02; Pearson r = 0.1, p = 0.02; **Fig.6C**). In other words, stronger circuit inhibition (more negative *c_1_*) corresponded to a weaker order bias (smaller *ε*).

**Figure 6.**
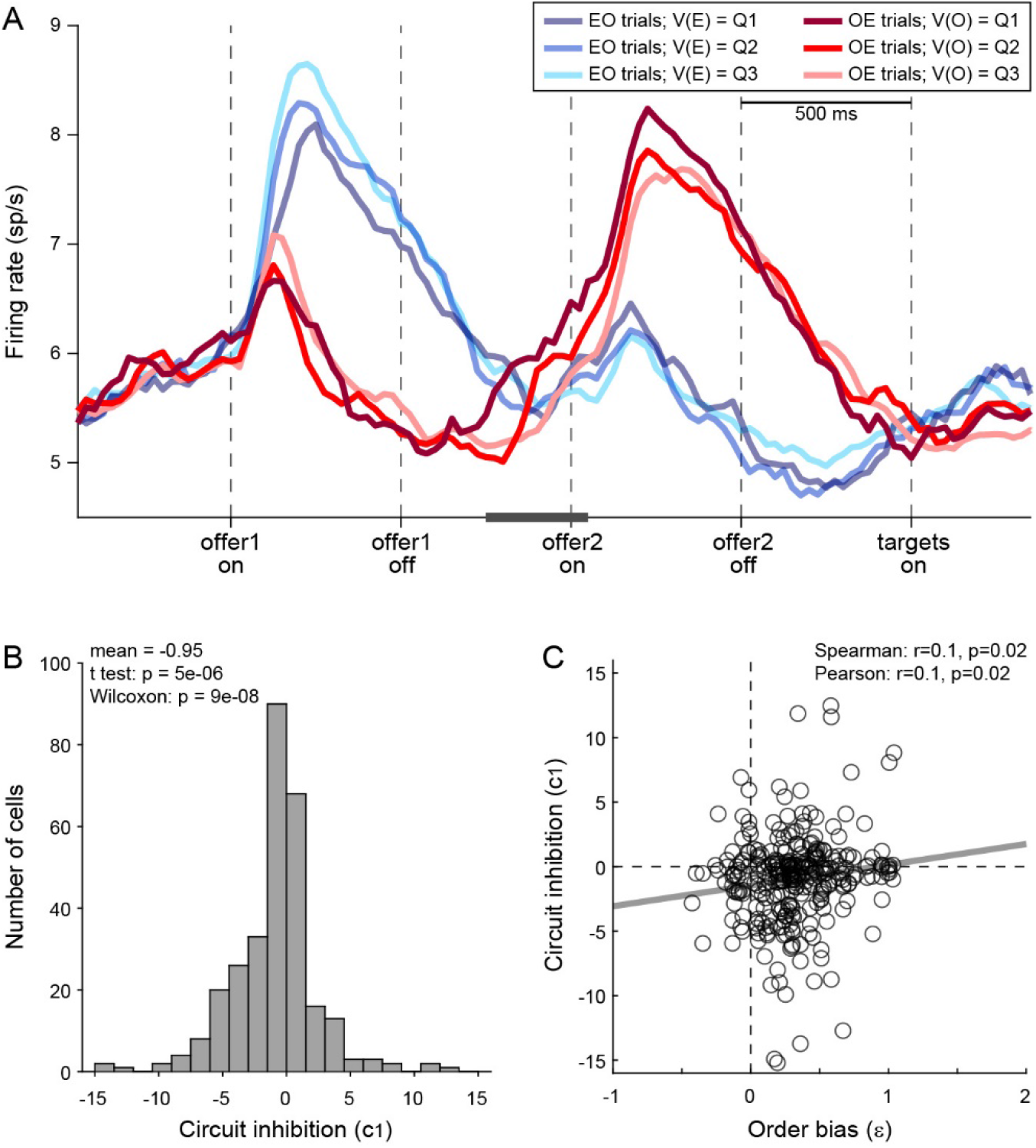
Order bias and circuit inhibition. **(A)** Circuit inhibition in *chosen juice* cells (primary data set, N = 160 cells). For each *chosen juice* cell E and O indicated the encoded juice and the other juice, respectively. We separated EO and OE trials, and divided each group of trials in tertiles based on the value of offer1. For EO trials, this corresponded to V(E); for OE trials, it corresponded to V(O). In this panel, Q1, Q2 and Q3 indicate low, medium and high values of offer1. In OE trials, shortly before offer2, the activity of *chosen juice* cells was negatively correlated with V(O) – a phenomenon termed circuit inhibition. For a quantitative analysis of circuit inhibition, we focused on 300 ms time window starting 250 ms before offer2 onset (black line). **(B)** Circuit inhibition for individual cells (N = 295 cells). For each *chosen juice* cell, we regressed the firing rate against the normalized V(O) (see **Methods**). The histogram illustrates the distribution of regression slopes (*c_1_*), which quantify circuit inhibition for individual cells. The effect was statistically significant across the population (mean = −0.95). **(C)** Correlation between order bias and circuit inhibition (N = 295 cells). Here the x-axis is the order bias (*ε*), the y-axis is circuit inhibition (regression slope *c_1_*) and each data point represents one cell. The two measures were significantly correlated across the population. Panel A includes only the primary data set; thus circuit inhibition shown here replicates previous findings (Ballesta and Padoa-Schioppa, 2019). Panels BC include both the primary and the additional data sets (see **Methods**). In panels BC, 47 cells were excluded from the analysis because measures of order bias (*ε*) or circuit inhibition (*c_1_*) were detected as outliers by the interquartile criterion. Including these cells in the analysis did not substantially alter the results. Statistical tests and exact p values are indicated in panels BC.

In conclusion, the order bias did not originate before or during valuation. Conversely, analysis of *chosen juice* cells and *chosen value* cells indicated that the order bias emerged during value comparison (decision stage).

### The preference bias emerged late in the trial (post-comparison)

When offers were presented sequentially (Task 2), both monkeys showed an additional preference bias that favored juice A and was more pronounced when the relative value of the two juices was larger (**Fig.3**). Our last series of analyses focused on the origins of this bias.

First, we inquired whether the preference bias emerged during valuation. If this was the case, one or both of the following should be true: (a) *offer value A* cells encoded higher values in Task 2 than in Task 1 and/or (b) *offer value B* cells encoded lower values in Task 2 than in Task 1. Furthermore, these putative effects should increase as a function of the relative value. To test these predictions, we examined the tuning functions of *offer value* cells. For each cell group *(offer value A, offer value B),* we pooled neurons with positive and negative encoding. For Task 1, we focused on the post-offer time window; for Task 2, we focused on post-offer1 and post-offer2 time windows, pooling trial types from both windows. Indicating with *b_0_* and *b_1_* the tuning intercept and tuning slope (**Eq.4**), we computed the difference in intercept *Δb_0_* = *b_0,Task2_* – *b_0,Task1_* and the difference in slope *Δb1* = *b_1,Task2_* – *b_1,Task1_* for each cell. We then examined the relation between these measures and the relative value *ρ* across the population, separately for each cell group. Contrary to the prediction, we did not find any correlation between neuronal measures (*Δb_0_*, *Δb_1_*) and the behavioral measure (*ρ*) for either *offer value A* or *offer value B* cells (**Fig.S3**). Thus the preference bias did not seem to emerge at the valuation stage.

We next examined *chosen value* cells. As discussed above, their activity provided a neuronal measure for the relative value (*ρ^neuronal^*), which reflected the internal subjective values of the juices emerging during value comparison. In principle, *ρ^neuronal^* might differ from the relative value derived from choices through the probit regression (*ρ^behavioral^*) because choices might be affected by systematic biases originating downstream of value comparison (**Fig.1**). In the light of this consideration, we examined the relation between the neuronal measure of relative value in Task 2 (*ρ^neuronal^_Task2_*, see **Methods**) and the behavioral measures obtained in the two tasks (*ρ^behavioral^_Task1_*, *ρ^behavioral^_Task2_*). We envisioned two possible scenarios (**Fig.7A**). In scenario 1, the preference bias reflected a difference in values across tasks. In other words, the subjective values of the juices in the two tasks were different and such that the relative value of juice A was higher in Task 2 than in Task 1. If so, *ρ^neuronal^_Task2_* should be statistically indistinguishable from *ρ^behavioral^_Task2_* and systematically larger than *ρ^behavioral^_Task1_*. In scenario 2, the subjective values of the juices were the same in both tasks and the preference bias reflected some neuronal process taking place downstream of value comparison. If so, *ρ^neuronal^_Task2_* should be statistically indistinguishable from *ρ^behavioral^_Task1_* and systematically smaller than *ρ^behavioral^_Task2_*.

**Figure 7.**
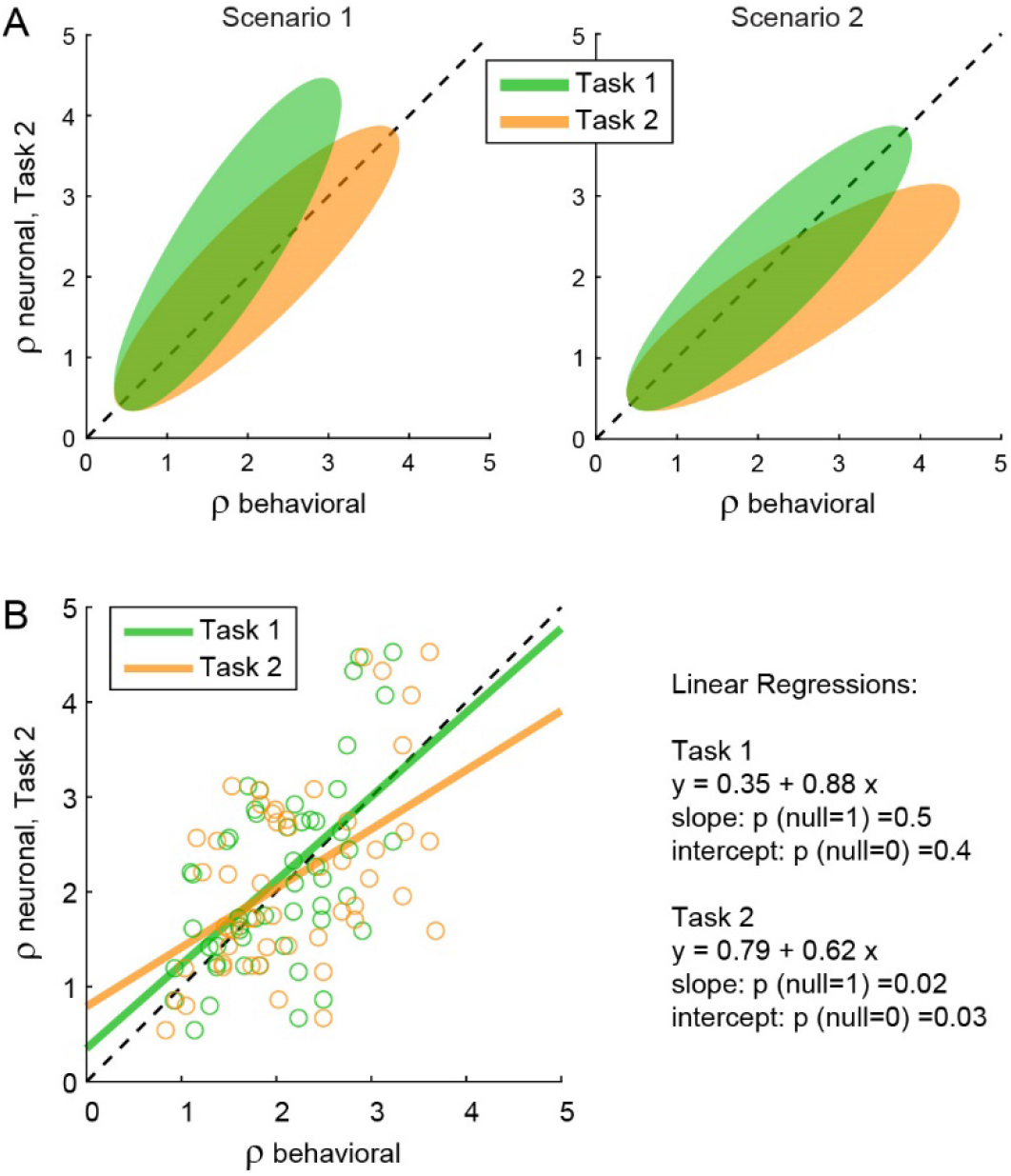
The preference bias does not reflect differences in the activity of *chosen value* cells. **(A)** Hypothetical scenarios. The two panels represent in cartoon format two possible scenarios envisioned at the outset of this analysis. In both panels, the x-axis represents behavioral measures from either Task 1 (green) or Task 2 (yellow); the y-axis represents the neuronal measure from Task 2. In scenario 1, the animal assigned higher relative value to juice A in Task 2. Thus, neuronal measures of relative value derived from the activity of *chosen value* cells in Task 2 (*ρ^neuronal^_Task2_*) would align with behavioral measures from the same task (*ρ^behavioral^_Task2_*) and be systematically higher than behavioral measures from Task 1 (*ρ^behavioral^_Task1_*). In scenario 2, the animal assigned the same relative values to the juices in both tasks. Thus, neuronal measures of relative value in Task 2 (*ρ^neuronal^_Task2_*) would be systematically lower than behavioral measures from the same task (*ρ^behavioral^_Task2_*) and would align with behavioral measures from Task 1 (*ρ^bebavioral^_Task1_*). **(B)** Empirical results (N = 52 cells). Neuronal measures derived from Task 2 (*ρ^neuronal^_Task2_*) are plotted against behavioral measures obtained in Task 1 (*ρ^behavioral^_Task1_*, green) or Task 2 (*ρ^behavioal^_Task2_*, yellow. Lines are from linear regressions. In essence, *ρ^neuronal^_Task2_* was statistically indistinguishable from *ρ^behavioral^_Task1_* and systematically lower than *ρ^behavioral^_Task2_*. Details on the statistics and exact p values are indicated in the figure. The analysis was restricted to 52 cells that had significant *θ_1A_*, *θ_1B_*, *θ_2A_* and *θ_2B_*. For this analysis, *ρ^neuronal^_Task2_* was taken as equal to *ρ^neuronal^_offer2_* (**Eq.10**). Other definitions provided similar results (data not shown).

The results of our analysis clearly conformed with scenario 2 (**Fig.7B**). For each *chosen value* cell, we computed *ρ^neuronal^_Task1_* in the post-offer time window and *ρ^neuronal^_Task2_* in the post-offer2 time window. Across the population, the two measures were statistically indistinguishable (p = 0.3, t test; not shown). We then regressed *ρ^neuronal^_Task2_* onto *ρ^behavioral^_Task1_*. The linear relation between these measures was statistically indistinguishable from identity. Separately, we regressed *ρ^neuronal^_Task2_* onto *ρ^behavioral^_Task2_*. In this case, the regression slope was significantly <1 (p = 0.02). This result is quite remarkable. It shows that the chosen value represented in the brain in Task 2 was equal to that inferred from choices in Task 1, and significantly different from that inferred from choices in Task 2. This fact implies that the preference bias was costly for the monkey, as it reduced the value obtained on average at the end of each trial.

In summary, the preference bias did not reflect differences in the values assigned to individual offers (offer values). Furthermore, insofar as the activity of *chosen value* cells reflects the decision process (**Fig.1**), the preference bias did not seem to emerge during value comparison. So how can one make sense of this behavioral phenomenon? At the cognitive level, the preference bias might be interpreted as due to the higher demands of Task 2. When the two saccade targets appeared on the monitor, information about values was no longer on display (**Fig.2B**). If at that point the animal had not finalized its decision, or if it had failed to retain in working memory the decision outcome, the animal might have selected the target associated with the better juice (juice A). Such bias would have been especially strong when the value difference between the two juices was large. In this view, the preference bias would reflect a “second thought” occurring after value comparison, in some trials.

To test this intuition, we turned to the activity of *chosen juice* cells. As noted above, in Task 2, the evolving decision was captured by the activity of these neurons recorded in OE trials immediately before and after offer2 onset (**Fig.8A**). More specifically, the state of the ongoing decision was captured by the distance between the two traces corresponding to the two possible choice outcomes (E chosen, O chosen). For any neuron, we quantified this distance with an ROC analysis, which provided a choice probability (CP). In essence, CP can be interpreted as the probability with which an ideal observed may guess the eventual choice outcome based on the activity of the cell. For each *chosen juice* cell, we computed the CP at different times in the trial. Across the population, mean(CP) exceeded chance level starting shortly before offer2, consistent with the above discussion on circuit inhibition. We then proceeded to investigate the origins of the preference bias.

**Figure 8.**
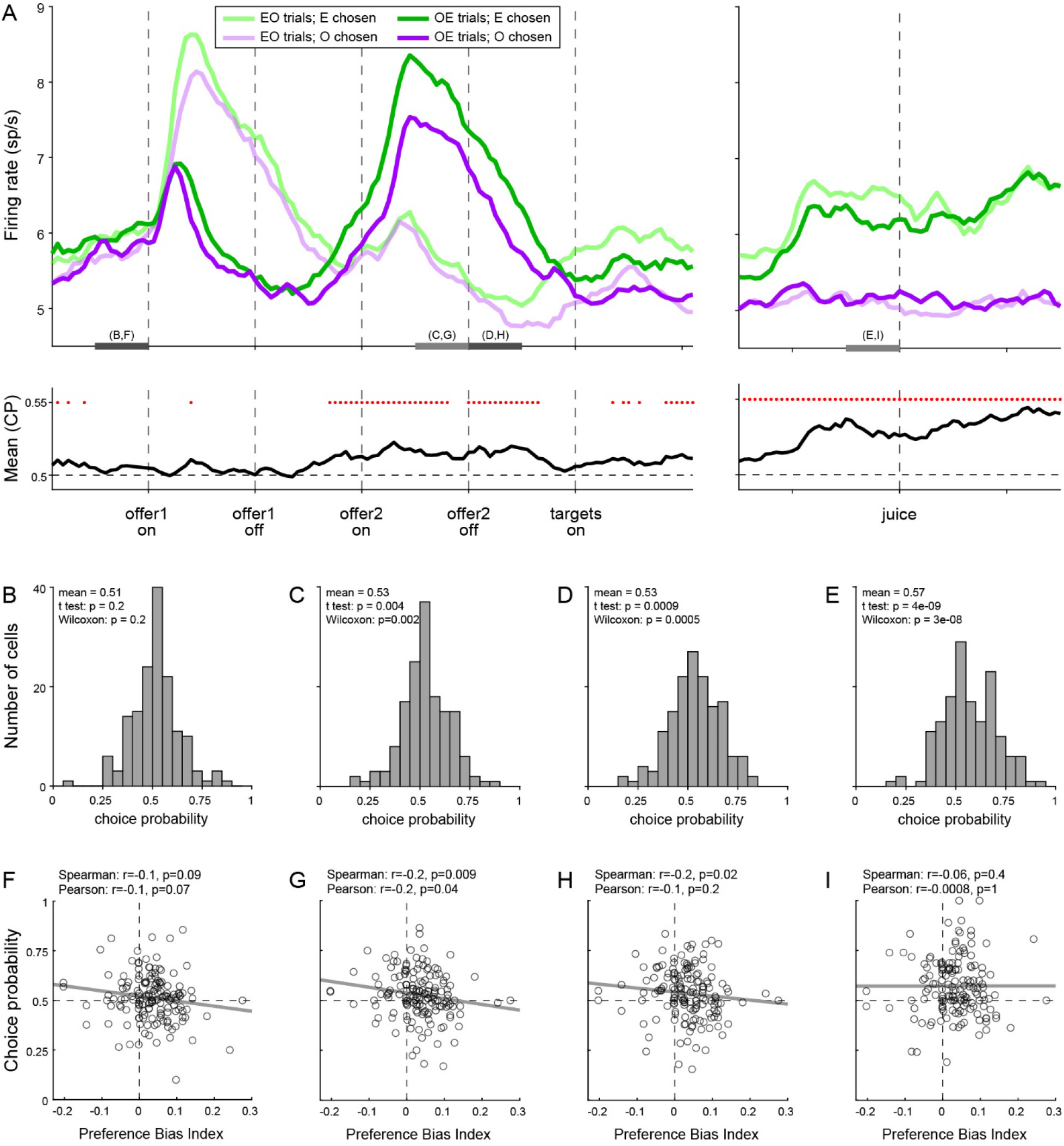
Preference bias and choice probability in *chosen juice* cells. **(A)** Profiles of activity and choice probability (N = 160 cells). On the top, separate traces are activity profiles for EO trials (dark colors) and OE trials (light colors), separately for E chosen (blue) and O chosen. On the bottom the trace is the mean(CP) computed for OE trials in 100 ms sliding time windows (25 ms steps). Red dots indicate that mean(CP) was significantly >0.5 (p<0.001; t test). Value comparison typically takes place shortly after the onset of offer2. **(B-E)** Distribution of CP in four 250 ms time windows. The time windows used for this analysis are indicated in panel A. **(F-I)** Correlation between CP and preference bias index. Each panel corresponds to the histogram immediately above it. CPs are plotted against the preference bias index (PBI), which quantifies the preference bias independently of the juice types. Each symbol represents one cell and the line is from a linear regression. CP and PBI were negatively correlated immediately before and after offer2 onset, but not later in the trial. This pattern suggests that the preference bias emerged late in the trial, when decisions were not finalized shortly after offer2 presentation.

We reasoned that, at the net of noise in measurements and cell-to-cell variability, CPs ultimately quantify the animal’s commitment to the eventual choice outcome. If the preference bias emerged late in the trial – perhaps after target presentation, if animals had not already finalized their decision – the intensity of the preference bias should be inversely related to the animals’ commitment to the eventual choice outcome measured earlier in the trial. In other words, there should be a negative correlation between the preference bias and CPs computed at the time when decisions normally take place (shortly before or after offer2 onset). Our analyses supported this prediction. To quantify the preference bias intensity independent of the juice pair, we defined the preference bias index PBI = 2 (*ρ_Task2_* – *ρ_Task1_*) / (*ρ_Task2_* + *ρ_Task1_*). We then focused on four 250 ms time windows before offer1 (control window), before and after offer2 onset, and before juice delivery (**Fig.8B-E**). Confirming our predictions, CP and PBI were significantly anti-correlated immediately before and during offer2 presentation, but not in the control time window or late in the trial (**Fig.8F-I**).

In conclusion, our results indicated that the preference bias did not emerge during valuation or during value comparison. Conversely, our results suggest that the preference bias emerged late in the trial, as a “second thought” process that guided choices when decisions were not finalized based on offer values alone. Notably, the preference bias effectively reduced the value monkeys obtained on average on any given trial.

## Discussion

Early economists proposed that choices between goods entail the computation and comparison of subjective values. However, the concept of value is somewhat slippery, because values relevant to choices cannot be measured behaviorally other than from choices themselves. This circularity problem hunted generations of scholars, dominating academic debates in the 19^th^ and 20^th^ century. In the end, neoclassic economic theory came to reject (cardinal) values and to rely only on (ordinal) preferences (Niehans, 1990; Samuelson, 1947). In other words, standard economics is agnostic as to whether subjective values are computed at all. The construction of standard economic theory was a historic success, but it came at a cost: the theory cannot explain a variety of biases observed in human choices (Camerer et al., 2003; Kahneman and Tversky, 2000; Lichtenstein and Slovic, 2006). In this perspective, neuroscience results showing that neuronal activity in multiple brain regions is linearly related to values defined behaviorally (Amemori and Graybiel, 2012; Cai et al., 2011; Cai and Padoa-Schioppa, 2012; Hosokawa et al., 2013; Jezzini and Padoa-Schioppa, 2020; Kable and Glimcher, 2007; Kim et al., 2008; Lak et al., 2014; Levy et al., 2010; Louie and Glimcher, 2010; Padoa-Schioppa and Assad, 2006; Pastor-Bernier et al., 2019; Plassmann et al., 2007; Shenhav and Greene, 2010), constitute a significant breakthrough. They validate the concept of value and effectively break the circularity surrounding it. Indeed, a neuronal population whose activity is reliably correlated with values measured from choices (behavioral values) may be used to derive independent measures of subjective values (neuronal values). In most circumstances, neuronal values and behavioral values should be (and are) indistinguishable. However, in specific choice contexts, the two measures might differ somewhat. When observed, such discrepancies indicate that choices are partly determined by processes that escape the maximization of offer values. If so, suitable analyses of neuronal activity may be used to assess the origins of particular choice biases.

These considerations motivated the analyses conducted in this study. In our experiments, monkeys chose between two juices offered simultaneously or sequentially. Choices under sequential offers were less accurate, biased in favor of the second offer (order bias), and biased in favor of the preferred juice (preference bias) (Shi et al., 2021). Earlier work had identified in OFC three groups of neurons encoding individual offer values, the chosen juice and the chosen value. Furthermore, earlier work indicated that these cell groups constitute the building blocks of a decision circuit (Padoa-Schioppa and Conen, 2017). In this view, *offer value* cells provide the primary input to a circuit formed by *chosen juice* cells and *chosen value* cells, where decisions are formed. Different cell groups in OFC may thus be associated with different computational stages: *offer value* cells instantiate the valuation stage; *chosen value* cells reflect values possibly modified by the decision process; and *chosen juice* cells capture the evolving commitment to a particular choice outcome. In this framework, we examined the activity of each cell group in relation to each behavioral phenomenon.

Our results may be summarized as follows. (1) Other things equal, neuronal signals encoding the offer values were weaker (smaller activity range) under sequential offers than under simultaneous offers. The reason for this discrepancy is unclear, but this neuronal effect was correlated with the difference in choice accuracy measured at the behavioral level. In other words, the drop in choice accuracy observed under sequential offers originated, at least partly, at the valuation stage. (2) The order bias did not correlate with any measure in the activity of *offer value* cells. However, the order bias was negatively correlated with circuit inhibition in *chosen juice* cells – a phenomenon seen as key to value comparison (Ballesta and Padoa-Schioppa, 2019). Furthermore, session-to-session fluctuations in the order bias correlated with fluctuations in the neuronal measure of relative value derived from *chosen value* cells. These findings indicate that the order bias emerged during value comparison. (3) The preference bias did not have any correlate in the activity of *offer value* cells or *chosen value* cells. Moreover, the preference bias was inversely related to a measure derived from *chosen juice* cells and quantifying the degree to which the decision was finalized when offer values are “normally” compared (i.e., following presentation of the second offer). These findings indicate that the preference bias emerged late in the trial.

Two findings are particularly relevant to the distinction between behavioral values and neuronal values. First, the activity of *offer value* cells did not present any difference associated with the presentation order or with the juice preference. Second, relative value derived from *chosen value* cells under sequential offers differed significantly from behavioral measures obtained in the same task, and were indistinguishable from behavioral measures obtained in the other task (simultaneous offers). Thus the order bias and the preference bias highlighted significant differences between neuronal and behavioral measures of value. These observations imply that the order bias and the preference bias emerged downstream of valuation. Importantly, they also imply that the two choice biases imposed a cost to the animals, in the sense that they reduced the (neuronal) value obtained on average in any given trial. Notably, it would be impossible to draw such conclusion based on choices alone.

To our knowledge, this is the first study to investigate the origins of choice biases building on the distinction between behavioral values and neuronal values. At the same time, some of our results are not unprecedented. Earlier work showed that human and animal choices are affected by a bias favoring, on any given trial, the same good chosen in the previous trial (Alos-Ferrer et al., 2016; Goodwin, 1977; Padoa-Schioppa, 2013; Schoemann and Scherbaum, 2019; Senftleben et al., 2021). The origins of this phenomenon, termed choice hysteresis, are hard to pinpoint based on behavioral evidence alone. However, previous analysis of neuronal activity in OFC revealed that choice hysteresis is not reflected in the encoding of offer values (Padoa-Schioppa, 2013). Conversely, choice hysteresis correlates with fluctuations in the baseline activity of *chosen juice* cells, which is partly influenced by the previous trial’s outcome. Thus, similar to the order bias, choice hysteresis appears to emerge at the decision stage.

To conclude, the past two decades have witnessed a lively interest for the neural underpinnings of choice behavior. In this effort, a significant breakthrough came from the adoption of behavioral paradigms inspired by the economics literature, in which subjective values derived from choices are used to interpret neural activity. Without renouncing this approach, here we took a further step, showing that the decision process sometimes falls short of selecting the maximum offer value, and that choices are sometimes affected by processes taking place downstream of value comparison. In other words, behavioral values and neuronal values sometimes differ. These results might seem uncontroversial, but they have deep implications for economic theory and beyond. Looking forward, the framework developed here, in which the computation and comparison of offer values are central, but choices can also be affected by other processes accessible through neuronal measures, may help understand the origins of other choice biases.

## Methods

All the experimental procedures adhered to the *NIH Guide for the Care and Use of Laboratory Animals* and were approved by the Institutional Animal Care and Use Committee (IACUC) at Washington University.

### Animal subjects, choice tasks and neuronal recordings

This study presents new analyses of published data (Shi et al., 2021). Experimental procedures for surgery, behavioral control and neuronal recordings have been described in detail. Briefly, two male monkeys *(Macaca mulatta;* monkey J, 10.0 kg, 8 years old; monkey G, 9.1 kg, 9 years old) participated in the study. Under general anesthesia, we implanted on each animal a head restraining device and an oval chamber (axes 50×30 mm) allowing bilateral access to OFC. During the experiments, monkeys sat in an electrically insulated environment with their head fixed and a computer monitor placed at 57 cm distance. The gaze direction was monitored at 1 kHz using an infrared video camera (Eyelink, SR Research). Behavioral tasks were controlled through custom written software (https://monkeylogic.nimh.nih.gov) (Hwang et al., 2019) based on Matlab (v2016a; MathWorks Inc).

In each session, the animal chose between two juices labeled A and B (A preferred) offered in variable amounts. Trials with two choice tasks, referred to as Task 1 and Task 2, were pseudo-randomly interleaved. In both tasks, offers were represented by sets of colored squares displayed on the monitor. For each offer, the color indicated the juice type and the number of squares indicated the quantity. Each trial began with the animal fixating a large dot. After 0.5 s, the initial fixation point changed to a small dot or a small cross; the new fixation point cued the animal to the choice task used in that trial. In Task 1 (**Fig.2A**), cue fixation (0.5 s) was followed by the simultaneous presentation of the two offers. After a randomly variable delay (1-1.5 s), the center fixation point disappeared and two saccade targets appeared near the offers (go signal). The animal indicated its choice with an eye movement. It maintained peripheral fixation for 0.75 s, after which the chosen juice was delivered. In Task 2 (**Fig.2B**), cue fixation (0.5 s) was followed by the presentation of one offer (0.5 s), an inter-offer delay (0.5 s), presentation of the other offer (0.5 s), and a wait period (0.5 s). Two colored saccade targets then appeared on the two sides of the fixation point. After a randomly variable delay (0.5-1 s), the center fixation point disappeared (go signal). The animal indicated its choice with a saccade, maintained peripheral fixation for 0.75 s, after which the chosen juice was delivered. Central and peripheral fixation were imposed within 4-6 and 5-7 degrees of visual angle, respectively. Aside from the initial cue, the choice tasks were nearly identical to those used in previous studies (Ballesta and Padoa-Schioppa, 2019; Padoa-Schioppa and Assad, 2006).

For any given trial, *q_A_* and *q_B_* indicate the quantities of juices A and B offered to the animal, respectively. An “offer type” was defined by two quantities [*q_A_ q_B_*]. On any given session, we used the same juices and the same sets of offer types for the two tasks. For Task 1, the spatial configuration of the offers varied randomly from trial to trial. For Task 2, the presentation order varied pseudo-randomly and was counterbalanced across trials for any offer type. The terms “offer1” and “offer2” indicated, respectively, the first and second offer, independently of the juice type and amount. Trials in which juice A was offered first and trials in which juice B was offered first were referred as “AB trials” and “BA trials”, respectively. The spatial location (left/right) of saccade targets varied randomly. The juice volume corresponding to one square (quantum) was set equal for the two choice tasks and remained constant within each session. It varied across sessions between 70 and 100 μl for both monkeys. The association between the initial cue (small dot, small cross) and the choice task varied across sessions in blocks. Across sessions, we used 12 different juices (and colors) and 45 different juice pairs. Based on a power analysis, in most sessions the number of trials for Task 2 was set equal to 1.5 times that for Task 1.

Neuronal recordings were guided by structural MRI scans (1 mm sections) obtained before and after surgery and targeted area 13m (Ongur and Price, 2000). We recorded from both hemispheres in both monkeys. Tungsten single electrodes (100 μm shank diameter; FHC) were advanced remotely using a custom-built motorized micro-drive. Typically, one motor advanced two electrodes placed 1 mm apart, and 1-2 such pairs of electrodes were advanced unilaterally or bilaterally in each session. Neural signals were amplified (gain: 10,000) band-pass filtered (300 Hz - 6 kHz; Lynx 8, Neuralynx), digitized (frequency: 40 kHz) and saved to disk (Power 1401, Cambridge Electronic Design). Spike sorting was performed off-line (Spike2, v6, Cambridge Electronic Design). Only cells that appeared well isolated and stable throughout the session were included in the analysis.

### Preliminary analyses

The present analyses build on the results of a previous study showing that both choice tasks engage the same groups of neurons in OFC (Shi et al., 2021). Here we briefly summarize those findings.

The original data set included 1,526 neurons (672 from monkey J, 854 from monkey G) recorded in 306 sessions (115 from monkey J, 191 from monkey G). In each session, choice patterns were analyzed using probit regressions as described in the main text (**Eq.1** and **Eq.2**). For Task 1 (simultaneous offers), the probit fit provided measures for the relative value *ρ_Task1_* and the sigmoid steepness *η_Task1_*. For Task 2 (sequential offers), the probit fit provided measures for the relative value *ρ_Task2_*, the sigmoid steepness *η_Task2_* and the order bias *ε*. For each neuron, trials from Task 1 and Task 2 were first analyzed separately using the procedures developed in previous studies (Ballesta and Padoa-Schioppa, 2019; Padoa-Schioppa and Assad, 2006). For Task 1, we defined four time windows: post-offer (0.5 s after offer onset), late-delay (0.5-1 s after offer onset), pre-juice (0.5 s before juice onset) and post-juice (0.5 s after juice onset). A “trial type” was defined by two offered quantities and a choice. For Task 2, we defined three time windows: post-offer1 (0.5 s after offer1 onset), post-offer2 (0.5 s after offer2 onset) and post-juice (0.5 s after juice onset). A “trial type” was defined by two offered quantities, their order and a choice. For each task, each trial type and each time window, we averaged spike counts across trials. A “neuronal response” was defined as the firing rate of one cell in one time window as a function of the trial type. Neuronal responses in each task were submitted to an ANOVA (factor: trial type). Neurons passing the p<0.01 criterion in ≥1 time window in either task were identified as “task-related” and included in subsequent analyses.

Following earlier work (Padoa-Schioppa, 2013), neurons in Task 1 were classified in one of four groups *offer value A*, *offer value B*, *chosen juice* or *chosen value*. Each variable could be encoded with positive or negative sign, leading to a total of 8 cell groups. Each neuronal response was regressed against each of the four variables. If the regression slope *b_1_* differed significantly from zero (p<0.05), the variable was said to “explain” the response. In this case, we set the signed *R^2^* as *sR^2^* = sign(*b_1_*) *R^2^;* if the variable did not explain the response, we set *sR^2^* = 0. After repeating the operation for each time window, we computed for each cell the *sum*(*sR^2^*) across time windows. Neurons explained by at least one variable in one time window, such that *sum*(*sR^2^*) ≠ 0, were said to be tuned; other neurons were labeled “untuned”. Tuned cells were assigned to the variable and sign providing the maximum |*sum*(*sR^2^*)|, where |·| indicates the absolute value. Thus indicating with “+” and “-” the sign of the encoding, each neuron was classified in one of 9 groups: *offer value A*+, *offer value A*-, *offer value B*+, *offer value B*-, *chosen juice A*, *chosen juice B*, *chosen value*+, *chosen value*- and *untuned*.

Neuronal classification in Task 2 followed the procedures described in a previous study (Ballesta and Padoa-Schioppa, 2019). Under sequential offers, neuronal responses in OFC were found to encode different variables defined in relation to the presentation order (AB or BA). Specifically, the vast majority of responses were explained by one of 11 variables including one binary variable capturing the presentation order (*AB | BA*), six variables representing individual offer values *(offer value A | AB, offer value A | BA, offer value B | AB, offer value B | BA, offer value 1,* and *offer value 2),* three variables capturing variants of the chosen value (*chosen value*, *chosen value A*, *chosen value B*) and a binary variable representing the binary choice outcome (*chosen juice*). Each of these variables could be encoded with a positive or negative sign. Most neurons encoded different variables in different time windows. In principle, considering 11 variables, 2 signs of the encoding and 3 time windows, neurons might present a very large number of variable patterns across time windows. However, the vast majority of neurons presented one of 8 patterns referred to as “sequences”. Classification proceeded as follows. For each cell and each time window, we regressed the neuronal response against each of the variables predicted by each sequence. If the regression slope *b_1_* differed significantly from zero (p<0.05), the variable was said to explain the response and we set the signed *R^2^* as *sR^2^* = sign(*b_1_*) *R^2^*; if the variable did not explain the response, we set *sR^2^* = 0. After repeating the operation for each time window, we computed for each cell the *sum*(*sR^2^*) across time windows for each of the 8 sequences. Neurons such that *sum*(*sR^2^*) ≠ 0 for at least one sequence were said to be tuned; other neurons were untuned. Tuned cells were assigned to the sequence that provided the maximum |*sum*(*sR^2^*)|. As a result, each neuron was classified in one of 9 groups: *seq #1*, *seq #2*, *seq #3*, *seq #4*, *seq #5*, *seq #6*, *seq #7*, *seq #8* and *untuned* (**Table S1**).

The results of the two classifications were compared using analyses for categorical data. In essence, we found a strong correspondence between the cell classes identified in the two choice tasks (Shi et al., 2021). Hence, we may refer to the different groups of cells using the standard nomenclature – *offer value, chosen juice* and *chosen value* – independently of the choice task. Based on this result, we proceeded with a comprehensive classification based on the activity recorded in both choice tasks. For each task-related cell, we calculated the *sum*(*sR^2^*) for the eight variables in Task 1 (*sum*(*sR^2^*)_*Task1*_) and eight sequences in Task 2 (*sum*(*sR^2^*)_*Task2*_) as described above. We then added the corresponding *sum*(*sR^2^*)_*Task1*_ and *sum*(*sR^2^*)_*Task2*_ to obtain the final *sum*(*sR^2^*)_*final*_. Neurons such that *sum*(*sR^2^*)_*final*_ ≠ 0 for at least one class were said to be tuned; other neurons were untuned. Tuned cells were assigned to the cell class that provided the maximum |*sum*(*sR^2^*)*_final_*|.

### Data sets

In some sessions, one or both choice patterns presented complete or quasi-complete separation – i.e., the animal split choices for <2 offer types in Task 1 and/or in Task 2. In these cases, the probit regression did not converge, the resulting steepness *η* was high and unstable, and the relative value was not unique. This issue affected the classification analyses described above only marginally, but for the present study it was critical that behavioral measures be accurate and precise. We thus restricted our analyses to stable sessions by imposing an interquartile criterion on the sigmoid steepness (Tukey, 1977). Defining IQR as the interquartile range, values below the first quartile minus 1.5*IQR or above the third quartile plus 1.5*IQR were identified as outliers and excluded. Thus our entire data set included 1,204 neurons (577 from monkey J, 627 from monkey G) recorded in 241 sessions (101 from monkey J, 140 from monkey G). In this population, the classification procedures identified 183 *offer value* cells, 160 *chosen juice* cells and 174 *chosen value* cells. These neurons constitute the primary data set for this study.

Most of our analyses compared choices and neuronal activity across tasks and were restricted to the primary data set. However, some analyses included only trials from Task 2 and quantified the effects due to the presentation order (AB vs. BA). In these analyses we included an additional data set recorded previously from the same two animals performing only Task 2 (Ballesta and Padoa-Schioppa, 2019). All the procedures for behavioral control and neuronal recording were essentially identical to those described above. Furthermore, behavioral analyses and inclusion criteria were identical to those used for the primary data set. The resulting data set included 1,205 neurons (414 from monkey J, 791 from monkey G) recorded in 196 sessions (51 from monkey J, 145 from monkey G). In this population, the classification procedures identified 243 *offer value* cells, 182 *chosen juice* cells and 187 *chosen value* cells. We refer to these neurons as the additional data set. Importantly, the order bias was also observed in these sessions (Ballesta and Padoa-Schioppa, 2019).

The interquartile criterion was also used to identify outliers in all the analyses conducted throughout this study. In practice, this criterion became relevant only for the analyses shown in **Fig.6** and **Fig.S2**, as indicated in the respective figure legends.

### Comparing tuning functions across choice tasks

Several analyses compared the tuning functions recorded in the two tasks (**Fig.4**, **Fig.S1-3**). Tuning functions were defined by the linear regression of the firing rate *r* onto the encoded variable *S*:

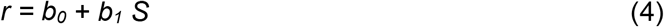

Regression coefficients *b_0_* and *b_1_* were referred to as tuning intercept and tuning slope, respectively. Positive and negative encoding corresponded to *b_1_*>0 and *b_1_*<0, respectively. We also defined the mean activity and the activity range as follows. Indicating with [*S_min_*, *S_max_*] the interval of variability for *S*, *ΔS* = *S_max_* – *S_min_* was the range of *S*. The mean activity was defined as *r_mean_* = *b_0_* + *b_1_ (S_max_+S_min_)/2*. The activity range was defined as *Δr* = |*b_1_ ΔS*|, where |·| indicates the absolute value.

For any neuronal response, the tuning was considered significant if *b_1_* differed significantly from zero (p<0.05) and if the sign of the encoding was consistent with the cell class (e.g., *b_1_*>0 for *offer value A* + cells). All the analyses comparing tuning functions across tasks were restricted to neuronal responses with significant tuning.

### Neuronal measures of relative value

Several analyses relied on neuronal measures for the relative value of the juices (*ρ^neuronal^*) derived from the activity of *chosen value* cells. In Task 1, these neurons encode the *chosen value* independently of the juice type. For each neuronal response, we performed a bilinear regression:

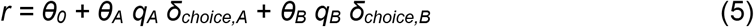

where *θ_0_*, *θ_A_* and *θ_B_* were the regression coefficients, *δ_choice,A_* = 1 if the animal chose juice A and 0 otherwise, and *δ_choice,B_* = 1 – *δ_choice,A_*. If the response encodes the *chosen value*, *θ_A_* should be proportional to the value of a quantum of juice A (uA), *θ_B_* should be proportional to the value of a quantum of juice B (uB), and the ratio *θ_A_*/*θ_B_* should equal the value ratio – i.e., the relative value of the two juices. We thus defined

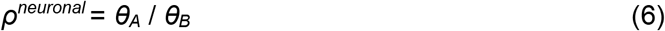

Previous studies showed that this measure is statistically indistinguishable from the behavioral measure *ρ^behavioral^* derived from the probit analysis of choice patterns (Padoa-Schioppa and Assad, 2006).

In Task 2, in the post-offer1 and post-offer2 time windows, *chosen value* cells encoded the value of the current offer, independent of the juice type (**Table S1**). For each neuron, we thus performed a bi-linear regression for each of the two time windows:

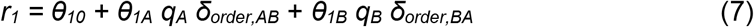

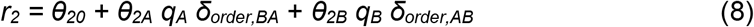

where *r_1_* and *r_2_* were their responses recorded in the post-offer1 and post-offer2 time windows, respectively, and *θ_10_*, *θ_1A_*, *θ_1B_*, *θ_20_*, *θ_2A_* and *θ_2B_* were regression coefficients. These coefficients provided four neuronal measures of relative value:

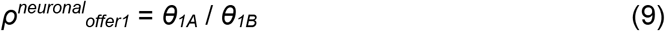

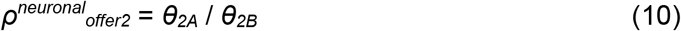

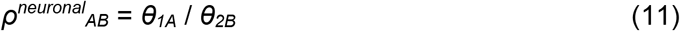

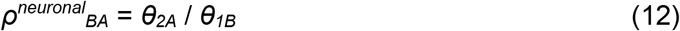

In essence, these four measures corresponded to the two time windows (post-offer1 and post-offer2) and to the two presentation orders (AB and BA). Importantly, all these measures were computed conditioned on *θ_1A_*, *θ_1B_*, *θ_2A_* and *θ_2B_* differing significantly from zero (p<0.05). The analyses illustrated in **Fig.5** and **Fig.7** were restricted to neurons satisfying this criterion.

In terms of notation, we often omit the superscript in *ρ^behavioral^* and we indicate behavioral measures simply as *ρ* (with the relevant subscripts). We use the superscript “behavioral” only when we explicitly compare behavioral and neuronal measures, for clarity. In contrast, for neuronal measures of relative value we always use the superscript “neuronal”.

### Activity profiles of chosen juice cells

To conduct population analyses, we pooled all *chosen juice* cells. The juice eliciting higher firing rates was labeled “E” (encoded) and other juice was labeled “O”. In Task 2, we thus referred to EO trials and OE trials, depending on the presentation order.

To illustrate the activity profiles of *chosen juice* cells in Task 2, we aligned spike trains at offer1 and, separately, at juice delivery. For each trial, the spike train was smoothed using a kernel that mimicked the post-synaptic potential by exerting influence only forward in time (decay time constant = 20 ms) (So and Stuphorn, 2010). In **Fig.6A** and **Fig.8A** we used moving averages of 100 ms with 25 ms steps for display purposes.

Under sequential offers, *chosen juice* cells encode different variables in different time windows (see **Table S1**). During offer1 and offer2 presentation, these cells encode in a binary way the juice type currently on display. Later, as the decision develops, these neurons gradually come to encode the binary choice outcome (i.e., the chosen juice). We previously showed that the activity of these neurons recorded in OE trials shortly before offer2 is inversely related to the value of offer1 (Ballesta and Padoa-Schioppa, 2019). This phenomenon, termed circuit inhibition, resembles the setting of a dynamic system’s initial conditions and is regarded as an integral part of the decision process (Ballesta and Padoa-Schioppa, 2019).

For a quantitative analysis of circuit inhibition, we focused on a 300 ms time window starting 250 ms before offer2 onset. We excluded forced choice trials, for which one of the two offers was null. For each neuron, we examined OE trials and we regressed the firing rates against the normalized value of offer1:

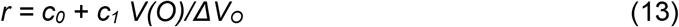

where *ΔV_O_* was the value range for juice O. The normalization allowed to pool neurons recorded with different value ranges. The regression slope *c_1_* quantified circuit inhibition for individual cells, and we studied this parameter at the population level.

The activity of *chosen juice* cells in OE trials captures the momentary state of the decision and thus the evolving commitment to a particular choice outcome. To quantify the momentary decision state, we conducted a receiver operating characteristic (ROC) analysis (Green and Swets, 1966) on the activity recorded during OE trials. This analysis was conducted on raw spike counts, without kernel smoothing, time averaging or baseline correction. We restricted the analysis to offer types for which the animal split choices between the two juices and we excluded trial types with <2 trials. For each offer type, we divided trials depending on the chosen juice (E or O) and we compared the two distributions. The ROC analysis provided an area under the curve (AUC). For each neuron, we averaged the AUC across offer types to obtain the overall choice probability (CP) (Kang and Maunsell, 2012). The ROC analysis was performed in 100 ms time windows shifted by 25 ms. We also conducted the same analysis on four 250 ms time windows, namely pre-offer1 (−250 to 0 ms from offer1 onset), late offer2 (−250 to 0 ms from offer1 offset), early wait (0 ms to 250 ms after offer2 offset) and pre-juice (−250 to 0 ms before juice delivery) (**Fig.8**). In **Fig.8B-I**, 6 cells were excluded because the Matlab function *perfcurve.m* failed to converge.

**Figure S1.**
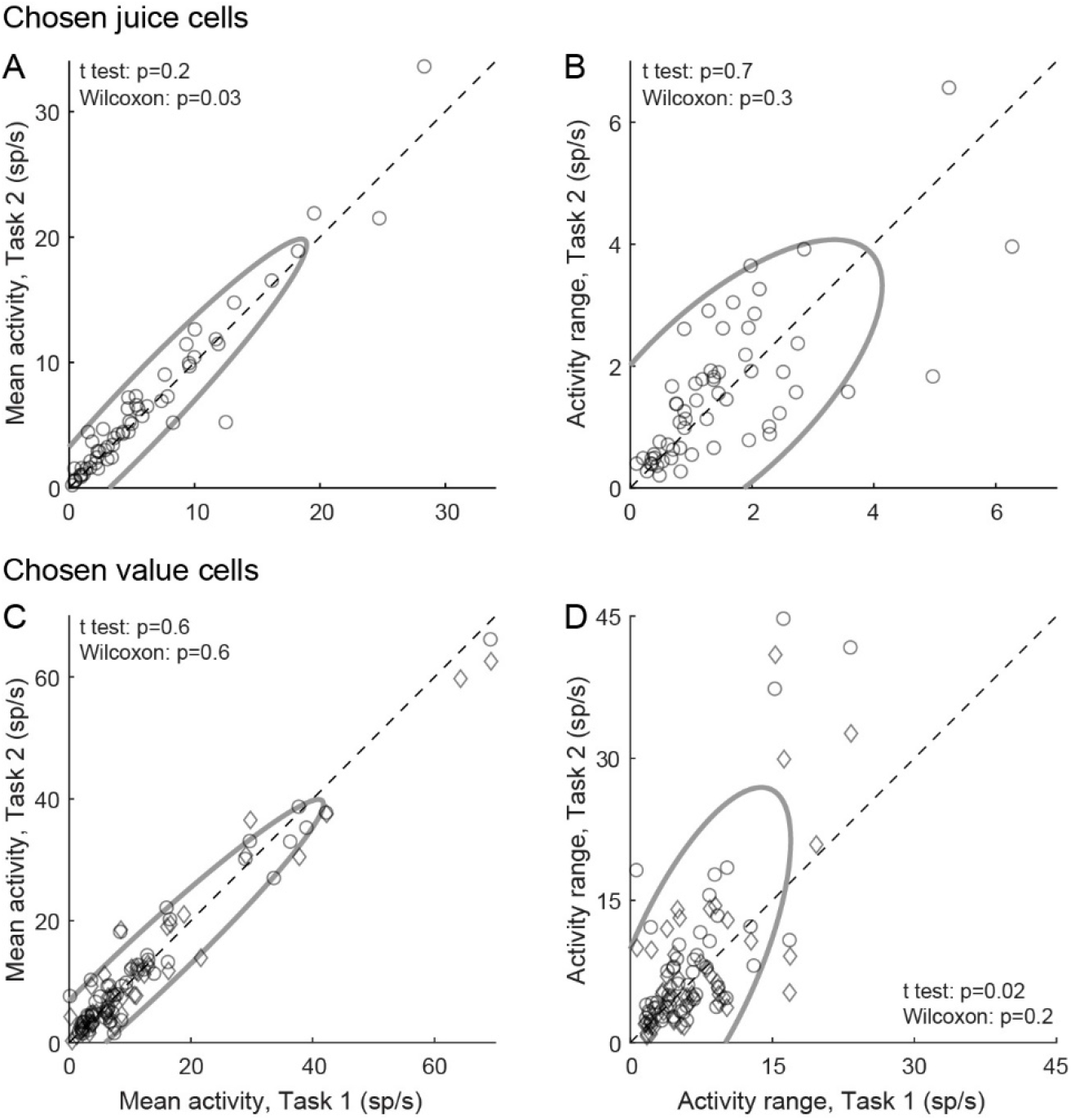
Comparing tuning functions across choice tasks. **(AB)** Chosen juice cells (N = 58). Same format as in **Fig.4AB**. For each cell, we examined the same time window (post-juice) in both tasks. Both the mean activity and the activity range were statistically indistinguishable across choice tasks. **(CD)** Chosen value cells (N = 104). For each cell, we examined one time window (post-offer) in Task 1 and two time windows (post-offer1 and post-offer2) in Task 2. Both the mean activity and the activity range were statistically indistinguishable across tasks. In panels B-G, legends report the results of statistical tests. For both cell groups, fluctuations in activity range were not correlated with fluctuations in choice variability across the population (in both analyses, |r| < 0.1, p > 0.4; not shown). Only cells presenting significant tuning in the relevant time windows were included in each panel (see **Methods**).

**Figure S2.**
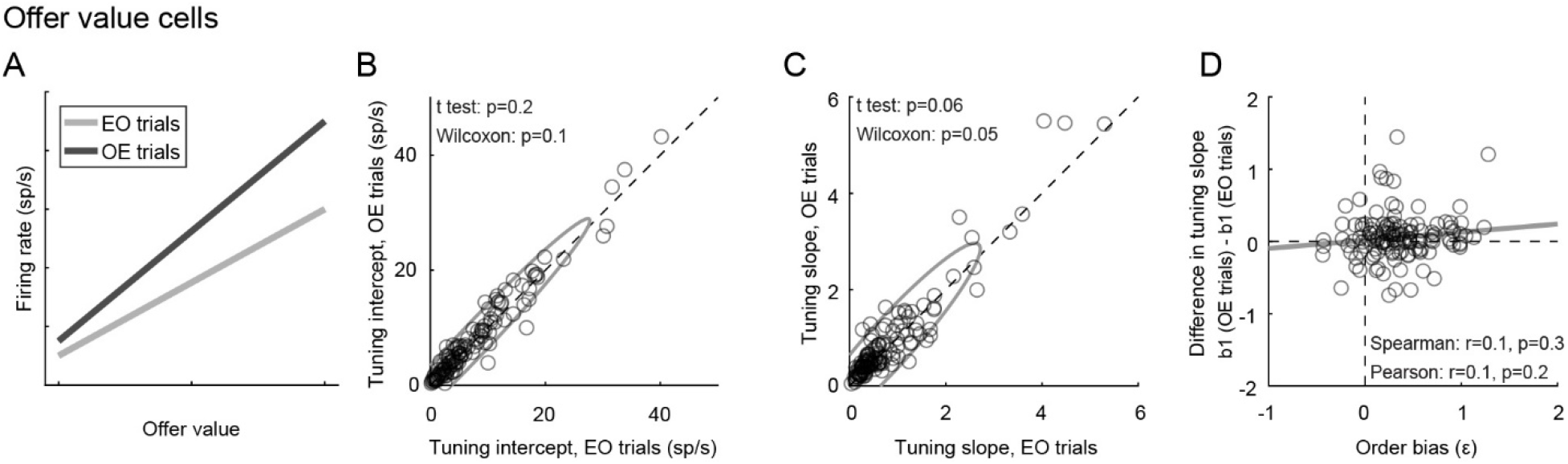
The order bias does not reflect differences in the tuning of *offer value* cells. **(A)** Rationale for the analysis. The two lines represent in cartoon format the hypothetical tuning functions of an *offer value* cell in the post-offer1 time window (EO trials) and in the post-offer2 time window (OE trials). The order bias would be explained if *offer value* cells encoded, other things equal, higher values in OE trials than in EO trials. This would be the case if the tuning intercept and/or the tuning slope were higher in OE trials, as depicted here. **(B)** Comparison of tuning intercepts. X- and y- axes represent the tuning intercept measured in post-offer1 (EO trials) and post-offer2 (OE trials) time windows, respectively. Each data point represents one cell. The two measures were statistically indistinguishable across the population. **(C)** Comparison of tuning slopes. Same format as panel B. The two measures were statistically indistinguishable across the population. **(D)** Lack of correlation between differences in tuning slope and order bias. Across the population, we did not find any correlation between the difference in tuning slope (y-axis) and the order bias. Exact p values are indicated in each panel. For this figure, we pooled neurons associated with A and B, and neurons with positive and negative encoding (N = 128 cells total). This analysis was restricted to cells significantly tuned in post-offer1 and post-offer2 time windows (Task 2). An additional 11 cells were removed because measures of order bias were detected as outliers by the interquartile criterion (see **Methods**). Including these cells in the analysis did not substantially alter the results. A similar analysis conducted on *chosen value* cells yielded similar negative results (not shown).

**Figure S3.**
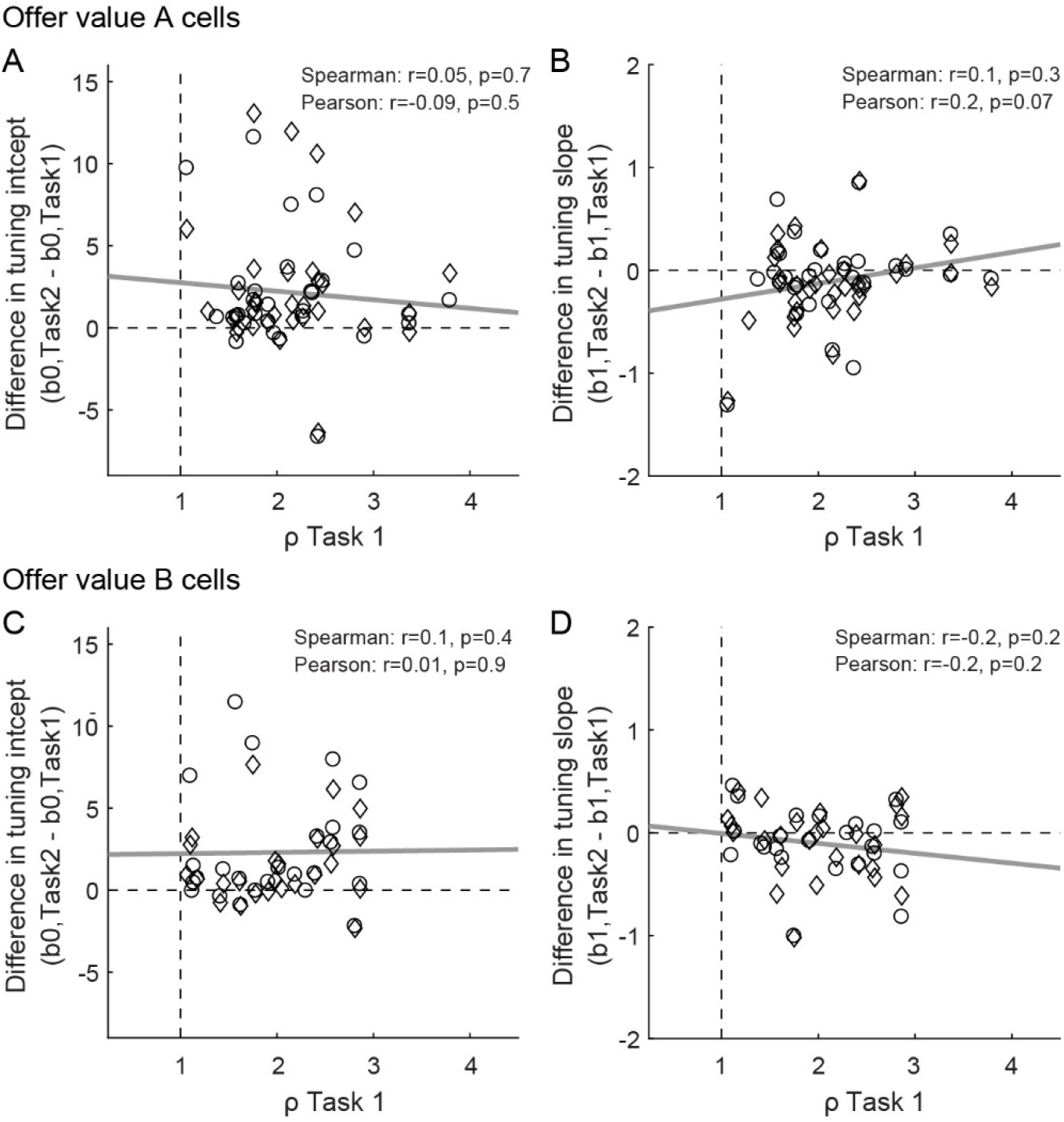
The preference bias does not reflect differences in the tuning of *offer value* cells. **(AB)** *Offer value A* cells (N = 63 cells). **(CD)** *Offer value B* cells (N = 51 cells). Panels A and C illustrate the relation between differences in tuning intercept (y-axis) and the relative value *ρ_Task1_* (x-axis); panels B and D illustrate the relation between differences in tuning slope (y-axis) and *ρ_Task1_* (x-axis). For each *offer value* cell, we examined one time window (post-offer) in Task 1 and two time windows (post-offer1 and post-offer2) in Task 2. In each panel, circles and diamonds refer to post-offer1 and post-offer2 time windows, respectively. Only cells presenting significant tuning in the relevant time windows were included in the analysis (see **Methods**). Exact p values are indicated in each panel and gray lines are from linear regressions. These analyses did not reveal any significant correlation.

**Table S1.**
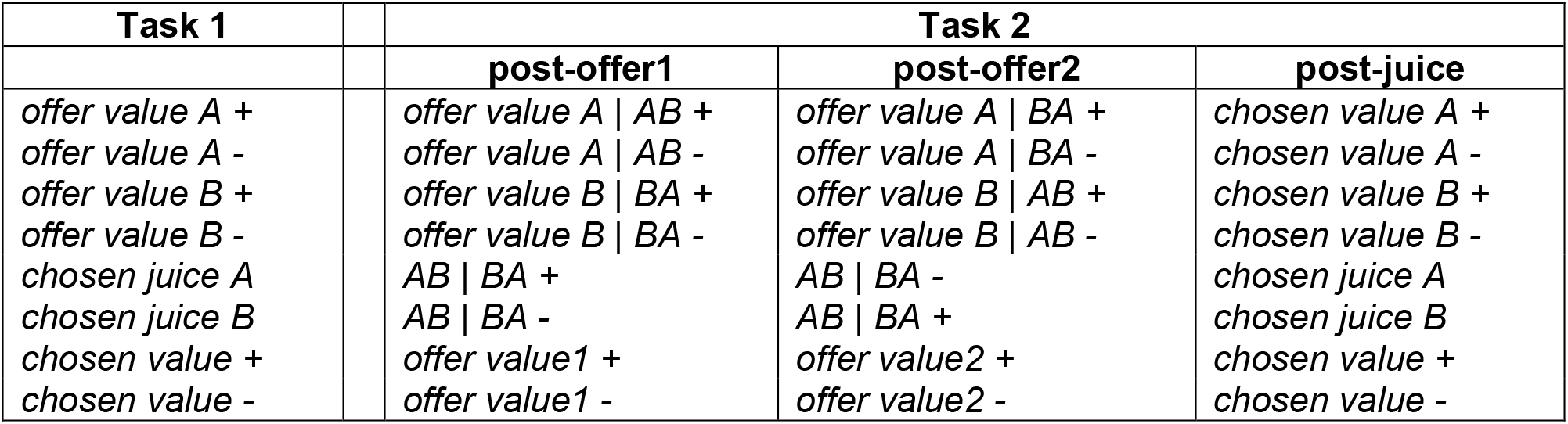
Neuronal encoding of decision variables in the two choice tasks. The table summarizes the results of a previous report (Shi et al., 2021). Under simultaneous offers, different groups of OFC neurons encode different decision variables, each with positive or negative sign (indicated here with + and −). In first approximation, each cell encodes the same variable across time windows. Under sequential offers, OFC neurons encode different variables in different time windows. However, the vast majority of them present one of 8 specific patterns of variables, referred to as variable “sequences” and detailed here. Furthermore, there is a clear correspondence between neurons encoding a particular variable in Task 1 and neurons encoding a particular sequence in Task 2. Hence, we can refer to different cell groups in OFC using the standard nomenclature originally defined for Task 1.

## Acknowledgments

We thank H. Schoknecht for help with animal training, L. Snyder for helpful discussions, and E. Bromberg-Martin, Z. Balewski, K. Conen, A. Livi, P. Natenzon, T. Ott, J. Tu and M. Zhang for comments on the manuscript. This research was supported by the National Institutes of Health (grant number R01-MH104494 to CPS) and by the McDonnell Center for Systems Neuroscience (pre-doctoral fellowship to WS).

## Conflict of interest

None

## Notes

### Competing Interest Statement

The authors have declared no competing interest.

